# Endocytic adaptor AP-2 maintains Purkinje cell function by balancing cerebellar parallel and climbing fiber synapses

**DOI:** 10.1101/2024.06.02.596459

**Authors:** M. Tolve, J. Tutas, E. Özer- Yildiz, I. Klein, E Koletsu, A. Petzold, F. Liebsch, Q. Silverman, M. Overhoff, G. Schwarz, T. Korotkova, S. Valtcheva, G. Gatto, NL Kononenko

## Abstract

The selective loss of cerebellar Purkinje cells is a hallmark of various neurodegenerative movement disorders, yet the precise mechanism driving their degeneration remains enigmatic. Here, we show that the endocytic adaptor protein complex 2 (AP-2) is essential for the survival of Purkinje cells. Employing a multidisciplinary approach encompassing mouse genetics, viral tracing, ex vivo calcium imaging, and kinematic analysis, we demonstrate that mice lacking the µ-subunit of AP-2 in cerebellar Purkinje cells exhibit early-onset ataxia associated with progressive Purkinje cell degeneration. Importantly, we uncover that synaptic input dysfunctions, characterized by a predominance of parallel fiber (PF) over climbing fiber (CF) synapses, precede Purkinje cell loss. Mechanistically, we find that AP-2 localizes to Purkinje cell dendrites, where it interacts with the PF synapse-enriched protein GRID2IP. The loss of AP-2 results in proteasome-dependent degradation of GRID2IP and accumulation of the glutamate δ2 receptor (GLURδ2) in distal Purkinje cell dendrites, leading to an excess of PF synapses while CF synapses are drastically reduced. The overrepresentation of PF synaptic input induces Purkinje cell hyperexcitation, which can be alleviated by enhancing synaptic glutamate clearance using the antibiotic ceftriaxone. Our findings demonstrate the critical role of AP-2 in preventing motor gait dysfunctions by regulating GRID2IP levels in Purkinje cells, thereby preserving the equilibrium of PF and CF synaptic inputs in a cell-autonomous manner.

## Introduction

Purkinje cells in the cerebellum play a vital role in motor learning and adaptation (Carey, 2024; Silva et al., 2024; Yang and Lisberger, 2014), and their dysfunction and/or degeneration is a hallmark of neurodegenerative movement disorders and animal models exhibiting ataxic symptoms. Despite extensive research, the precise mechanisms governing Purkinje cell survival remain elusive. The deterioration of Purkinje cell function is frequently accompanied by changes at the glutamatergic synapses. Several forms of spinocerebellar ataxia (SCA) can be caused by defects in glutamatergic transmission, including alterations at the two primary excitatory inputs to Purkinje cells: climbing fiber (CF) synapses, originating from the inferior olive, and parallel fiber (PF) synapses, composed of the axons of granule cells (Carey, 2011). In several SCA mouse models, the CFs are impaired in their development and/or connectivity (Anton et al., 2011; Barnes et al., 2011; Blake et al., 2013; Furrer et al., 2012; Smeets and Verbeek, 2016; Smeets et al., 2015). Interestingly, CF loss has been observed to precede Purkinje cell degeneration in some cases, indicating a potential role in disease progression (Smeets et al., 2015). The loss of CFs in SCAs is frequently accompanied by the strengthening of PF synapses, a process known as heterosynaptic competition (Hashimoto and Kano, 2013; Smeets et al., 2021; Sotelo et al., 1975). Given that CFs and PFs innervate distinct regions of the dendritic arbours of Purkinje cells – with CFs populating the proximal dendritic tree and PFs synapsing on the distal dendrites - the loss of one fiber type typically leads to an increase in the other (Miyazaki et al., 2012; Miyazaki et al., 2010; Taisuke et al., 2004). Maintaining the delicate balance between CF and PF synaptic inputs is crucial for synaptic plasticity and cerebellar gain control (Ohtsuki et al., 2009). However, the precise mechanisms governing the maintenance of this balance and its relationship to Purkinje cell degeneration remain poorly understood. A particularly intriguing and yet underexplored question is whether the balance of CF and PF synapses is regulated in a Purkinje cell-autonomous fashion.

A variety of signaling cascades at the plasma membrane, including IGF-1 and TrkB receptor signaling (Bosman et al., 2006; Kakizawa et al., 2003), ion channels (Cook et al., 2022; Hashimoto et al., 2011; Ichikawa et al., 2002; Kano et al., 1997; Miyazaki et al., 2010), and cell adhesion molecules (Zhang et al., 2015) are proposed to regulate heterosynaptic competition between CF and PF synapses. Here we investigate the role of the endocytic adaptor protein complex-2 (AP-2) (Conner and Schmid, 2003; Schmid and McMahon, 2007) in CF/PF synapse formation. AP-2 is a major orchestrator of clathrin-mediated endocytosis (CME) at the plasma membrane (Camblor-Perujo and Kononenko, 2022; Kononenko and Haucke, 2015; Robinson, 2004). AP-2 is comprised of α, β, μ, and σ subunits (Dittman and Ryan, 2009), and a mutation in its core μ subunit has been recently associated with developmental encephalopathy with ataxia (Helbig et al., 2019). While the brain-specific role of AP-2 has been extensively studied in cortical and hippocampal neurons - where it regulates synaptic vesicle biogenesis (Gu et al., 2013; Gu et al., 2008; Kim and Ryan, 2009; Kononenko et al., 2014), TrkB and BACE1 trafficking (Andres-Alonso et al., 2019; Bera et al., 2020; Kononenko et al., 2017), autophagy (Kononenko et al., 2017), and centrosome integrity (Camblor-Perujo et al., 2024) - its loss-of-function in the cerebellum and specifically in Purkinje cells, remains unexplored.

Here, we report that AP-2 regulates cerebellar function by controlling synaptic rewiring of PF and CF synapses in a cell-autonomous manner. Mice lacking the AP-2µ subunit in Purkinje cells exhibit early and severe gait disturbances along with progressive Purkinje cell degeneration. Notably, an increased number of dendritic spines and PF inputs precede Purkinje cell loss, while CFs are significantly reduced. Mechanistically, we demonstrate that AP-2 localizes to Purkinje cell dendrites, where it interacts with the PF synapse-enriched protein GRID2IP. The loss of AP-2 results in the proteasome-dependent degradation of GRID2IP and the accumulation of the glutamate δ2 receptor (GLURδ2) in distal Purkinje cell dendrites, leading to an excess of PF synapses. Remarkably, the overrepresentation of PFs induces Purkinje cell hyperexcitation, which can be alleviated by enhancing synaptic glutamate clearance with the antibiotic ceftriaxone. Our results highlight a novel role for the endocytic adaptor AP-2 in preventing motor gait dysfunction by regulating the dendritic levels of GRID2IP in Purkinje cells, thereby maintaining the balance of PF and CF synapses in the cerebellum.

## Results

### Mice lacking the endocytic adaptor AP-2 in Purkinje cells exhibit severe and early ataxia

To elucidate the precise role of AP-2 in cerebellar function, we developed a new mouse model in which the deletion of its central μ subunit is specifically driven in cerebellar Purkinje cells. To this end, we crossed previously published mice with floxed exons 2 and 3 of the *Ap2m1* gene (Kononenko et al., 2017) with mice expressing Cre-recombinase under the Purkinje cell-specific L7/*Pcp2*-promoter, whose activity is first observed around embryonic day 6 and is fully established 2 weeks after birth (Barski et al., 2000) (**Fig. 1A)**. Deletion of the μ-subunit destabilizes the AP-2 complex, leading to subsequent degradation of its α-subunit (Kononenko et al., 2014). Successful recombination was demonstrated by the expression of the Ai9-tdTomato reporter in WT mice and in mice lacking AP-2 in Purkinje cells (henceforth defined as conditional AP-2 cKO mice) (**Fig. S1A, see also Fig. S3D**) and by significant downregulation of the AP-2 complex in cerebellar lysates probed with an antibody against its α-subunit (**Fig. 1B,C**). CME function was perturbed upon loss of AP-2 in Purkinje cells, as evident by the asbcence of the characteristic clathrin foci in Purkinje cell dendrites (**Fig. S1B**), and by the inhibition of transferrin uptake in acute cerebellar slices from AP-2 cKO mice (**Fig. S1C,D**). The resulting offsprings were viable but showed a slightly reduced weight at 2 months of age (**Fig. S1E,F**) that was mainly due to the reduction of fat mass (**Fig. S1G**). To test whether AP-2 is required in Purkinje cells to mediate their function in motor coordination, we subjected the WT (i.e. *Ap2m1*^wt/wt^; L7*^Cre^*) and AP-2 cKO (i.e. *Ap2m1^fl/fl^*; L7*^Cre^*) mice to classical behavioural tasks, including the accelerating rotarod, treadmill, and self-paced locomotion on narrow beams (Miterko and Lin, 2021; Sathyamurthy et al., 2020). We found that 2-month-old AP-2 cKO mice were severely impaired in their ability to stay on the rotarod (**Fig. 1D**) and were not able to ambulate on a treadmill at increased speed as tested with DigiGait (**Fig. 1E,F, Videos S1,2**). Of note, we did not observe any locomotor problems in mice younger than 6-7 weeks (data not shown). Moreover, when challenged with beams of different widths, AP-2 cKO mice were able to cross the 25 mm-wide beams; however, they slipped significantly more often compared to their littermates (**Fig. 1G,H, Video S3,4**). Strikingly, these mice had extreme difficulty crossing the 12-mm beam and rarely managed to walk on a 5-mm beam (**Fig. 1H, Video S5-8**). To gain more insight into why AP-2 cKO mice are impaired in their ability to walk on the beam, we analyzed the kinematics of limb movements within each step cycle using AutoGaitA (Hosseini et al., 2024) (**Fig. S2A, Fig. 1I,J**). Alterations in limb kinematics were already visible on the 25 mm beam, with AP-2 cKO mice displaying reduced excursion of the ankle and knee joints, as well as a lowered iliac crest (**Fig. 1K,L, Fig. S2B,D,F,H,J**). As mice were challenged on a narrower beam (12 mm), these difference in kinematics became more pronounced, with the proximal joints (hip and iliac crest), the distal joints (ankle and knee), and the hindpaw being held closer to the beam surface (**Fig. 1M,N, Fig. S2C,E,G,I,K**). Furthermore, AP-2 cKO mice also displayed impaired intralimb joint coordination. By examining the angular changes across genotypes during the step cycle, we found that the dynamics of knee and ankle angles were massively altered in AP-2µ cKO mice (**Fig. S2L-Q**), while the hip angle, although showing a higher degree of variability, was similar to controls (**Fig. S2R,S**). Finally, the speed at which the paw and the ankle moved during the swing was greatly reduced in AP-2 cKO mice on both the 25- and 12-mm beams (**Fig. 1O-R**). Taken together, our data suggest that 2-month-old mice lacking the endocytic adaptor AP-2 in Purkinje cells exhibit reduced locomotor ability that is likely due to severe lower limb incoordination, a phenotype strongly reminiscent of ataxic gait in patients with cerebellar dysfunction (Ilg et al., 2007; Ilg and Timmann, 2013).

**Figure 1.**
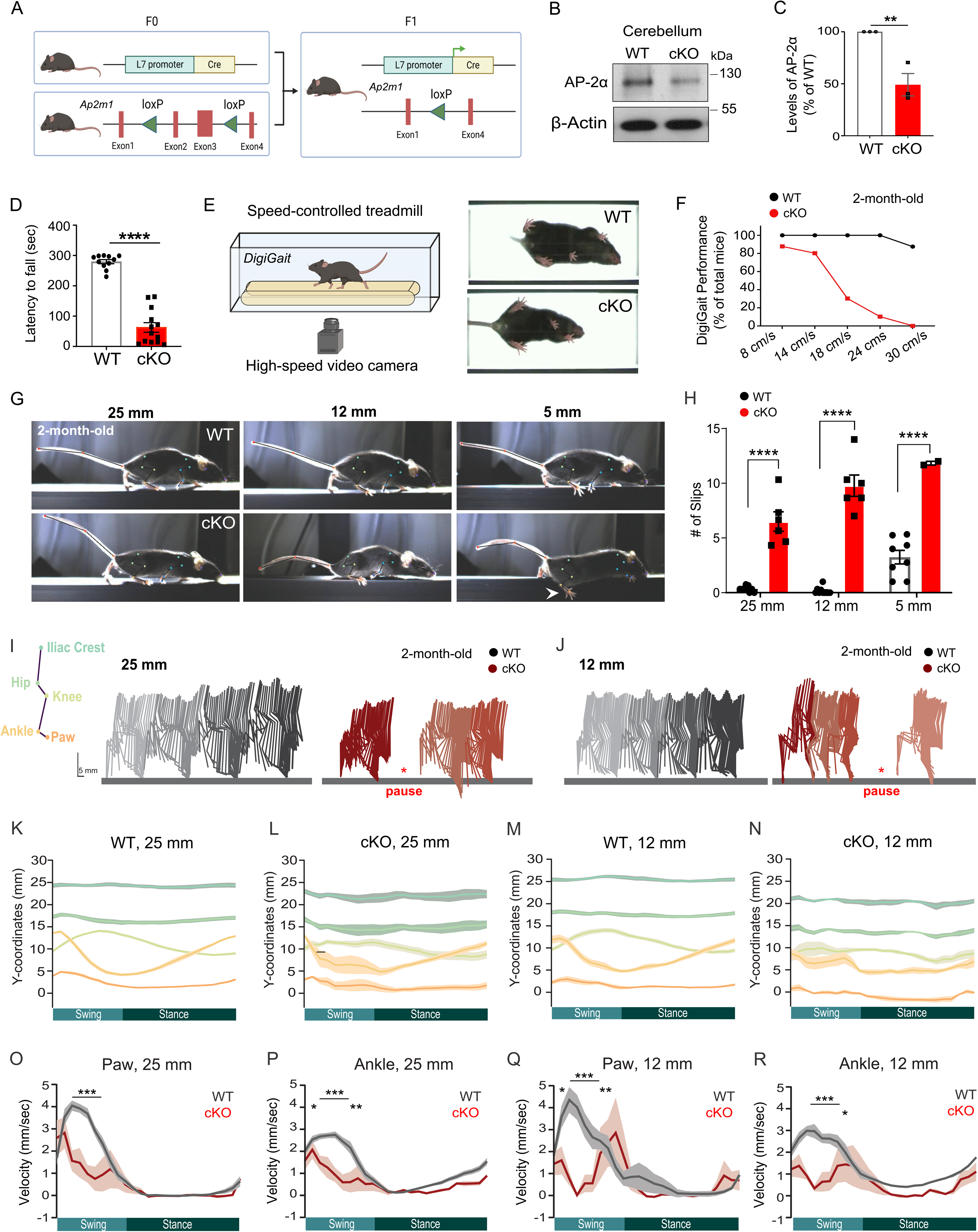
Loss of AP-2 in cerebellar Purkinje cells causes early ataxia in mice. **(A)** Schematic representation of the generation of AP-2 cKO mice. Conditional inactivation of the μ subunit of the AP-2 complex in cerebellar Purkinje cells was achieved by crossing *L7^Cre/+^* mice with *AP-2*_μ_*^flox/flox^* mice. **(B, C)** Representative immunoblot (B) and immunoblot analysis (C) of AP-2 levels in cerebellar lysates from 2-month-old WT and AP-2 cKO mice. AP-2, probed using antibody against its α subunit, is significantly reduced in cKO mice compared to WT set to 100. Each dot represents one mouse (N=3 for each genotype). Statistical significance was determined by one-tailed unpaired t-test (cKO: 49.08 ± 10.75 %; p=0.0045). **(D)** Rotarod performance of 2-month-old WT and AP-2 cKO mice. Latency to fall was significantly reduced in AP-2 cKO mice compared to WT controls. Each dot represents one mouse (N=11 for WT; N=13 for cKO). Statistical significance was determined by unpaired two-tailed Student’s t-test (WT: 279.6 sec ± 6.907 sec, cKO: 62.69 sec ± 15.93 sec; p<0.0001). **(E)** Ambulation of mice on a translucent treadmill (DigiGait) at controlled speeds (from 8 to 30 cm/s) to assess gait dynamics. Example images of ventral plane videography of 2-month-old WT and AP-2 cKO mice on the DigiGait treadmill. **(F)** DigiGait performance indicating the percentage of 2-month-old WT and AP-2 cKO mice able to ambulate on a treadmill at the indicated speeds (8cm/s: WT N=3, cKO N=8; 14 to 30 cm/s: WT N=6, cKO N= 10). **(G)** Example images of WT and AP-2 cKO mice crossing beams of different widths (25 mm, 12 mm and 5 mm). The white arrow shows the slip of an AP-2 cKO mouse on a 5 mm long bar. **(H)** Quantification of number of slips in 2 months-old WT and AP-2 cKO mice crossing beams of different widths. Each dot represents one mouse (25 and 12 mm: N=9 for WT, N=6 for cKO; 5 mm: N=9 for WT, N=2 for cKO). Statistical significance was determined by two-way ANOVA followed by Šidák multiple comparison test (25 mm: WT 0.34 ± 0.084, cKO 6.5 ± 0.90; p<0.0001. 12 mm: WT 0.16 ± 0.10, cKO 9.8 ± 0.97; p<0.0001. 5 mm: WT 3.2 ± 0.62, cKO 12 ± 0.17; p<0.0001). Note the drop out of cKO animals on 5 mm beam, indicating their inability to ambulate on a narrow path. **(I,J)** Stick diagrams representing kinematics of hindlimb movements within each step cycle in WT and AP-2 cKO mice crossing a 25 mm (I) or a 12 mm (J) beam. To be noted that in AP-2 cKO mice we could only analyse steps on the 25 mm and, partially, on the 12 mm beam, due to the numerous slips and falls. **(K-N)** Line graphs showing the variation in height (y-coordinates) of the iliac crest, hip, knee, ankle and hindpaw during a normalized step cycle in WT and AP-2 cKO mice walking on a 25mm- (K,L) or 12 mm-wide beam (M,N). Analysis of joint positioning during step cycle revealed significant changes in limb kinematics of AP-2 cKO mice already on the 25 mm beam (25 and 12 mm: N=9 for WT, N=6 for cKO). See Fig. S2B-K for a detailed two-way ANOVA-based pair-wise comparison of the kinematics for each joint in WT and AP_2 cKO mice. **(O,P)** Line graphs showing the variation in hindpaw (O) and ankle (P) velocities during a normalized step cycle in WT and AP-2 cKO mice walking on the 25mm beam. The velocity of paw (O) and ankle (P) was significantly altered during swing phase in AP-2 cKO mice compared to WT (N=9 for WT, N=6 for cKO). Statistical significance was determined by two-way ANOVA followed by Šidák multiple comparison. Statistical significance was determined by two-way ANOVA followed by Šidák multiple comparison between joints in WT and AP-2 cKO mice (see Table S4 for detailed results of two-way ANOVA multiple comparison). **(Q,R)** Line graphs showing the variation in hindpaw (Q) and ankle (R) velocities during a normalized step cycle in WT and AP-2 cKO mice walking on a 12 mm beam. The velocity of paw (Q) and ankle (R) was significantly altered during swing phase in AP-2 cKO mice compared to WT (N=9 for WT, N=6 for cKO). Statistical significance was determined by two-way ANOVA followed by Šídák multiple comparison. Data is presented as mean±SEM. Data presented in Fig. 1K-R show mean as a filled dark line, and SEM as the shaded area around it. * p ≤0.05; ** p≤0.01; *** p≤0.001; **** p≤0.0001.

### Loss of Purkinje cells in AP-2 cKO mice is preceded by their increased spine density

Ataxic gait in patients is usually accompanied by cerebellar atrophy (Hannoun and Hourani, 2022). Conversely, the loss of Purkinje cells, the major output of the cerebellar cortex, causes ataxia in mice (Hoxha et al., 2018). To further elucidate the mechanism of severe gait dysfunction in AP-2 cKO mice, we analyzed cerebellar morphology and Purkinje cell numbers in 2-month-old WT and AP-2 cKO mice. Immunohistochemical analysis of the AP-2 cKO cerebellum using the calbindin antibody to label Purkinje cells revealed that, although the gross morphology of the cerebellum was unaltered, the deletion of AP-2 resulted in a strong reduction in the number of Purkinje cells (**Fig. 2A**), a phenotype that was also observed in cerebellar slices from AP-2 cKO mice analyzed by Nissl staining (**Fig. 2B-D, Fig. S3A-C**). The decrease in Purkinje cell number was not due to developmental effects, as Purkinje cells were still present in the 1-month-old AP-2 cKO cerebellum (**Fig.2B**), and was progressive, with almost all AP-2-deficient Purkinje cells being lost by 3 months of age (**Fig. 2D, Fig. S3D**).

**Figure 2.**
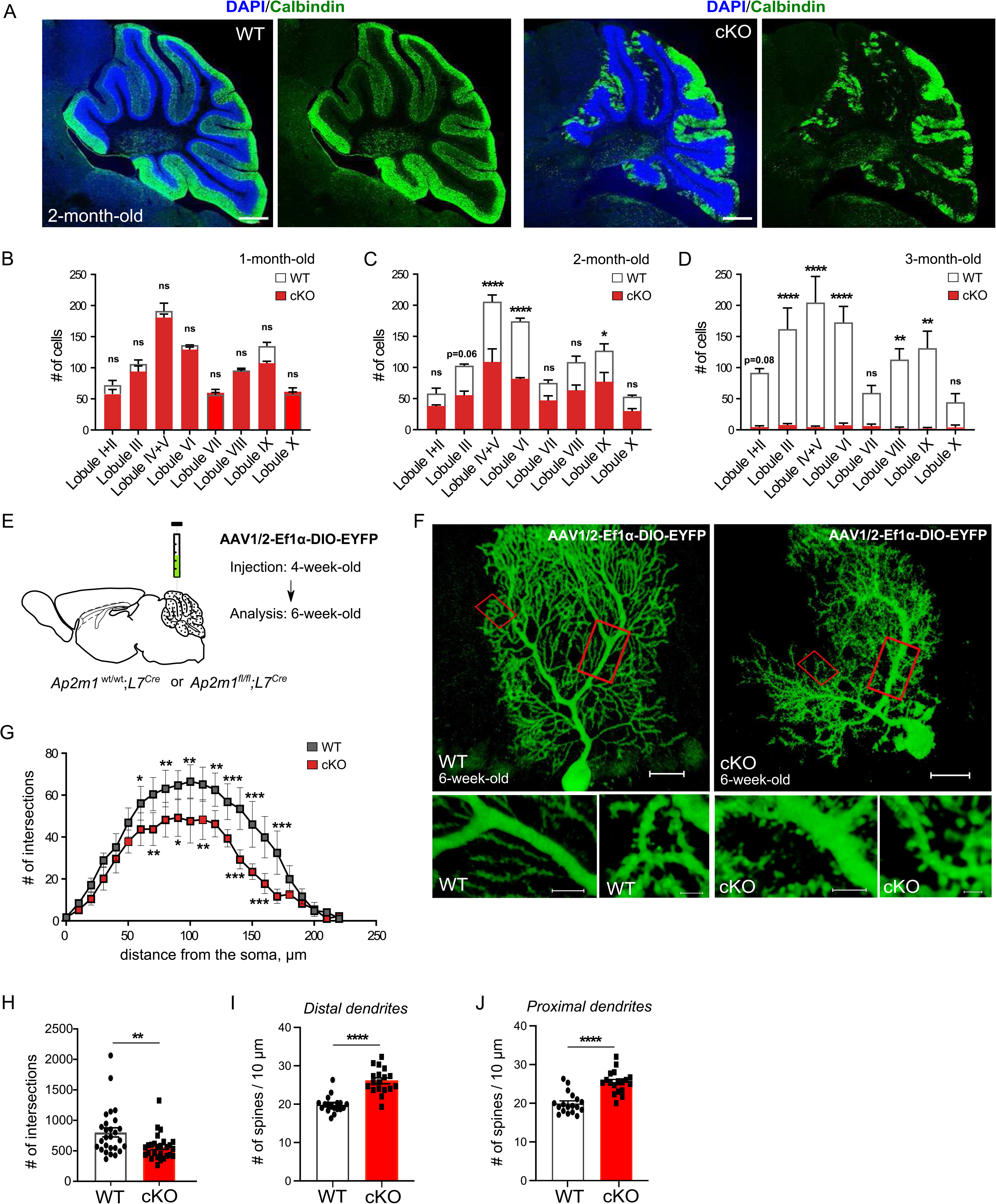
Progressive degeneration of AP-2 cKO Purkinje cells is preceded by their increased spine density. **(A)** Representative confocal images of WT and AP-2 cKO cerebellum at 2 months of age immunostained with an antibody against Calbindin to label Purkinje cells (green) and DAPI to label nuclei (blue). Scale bar: 500µm. **(B-D)** Quantification of Purkinje cell number per cerebellar lobule based on Nissl staining (see Fig. S3A-C) in 1 month- (B), 2 month- (C) and 3 month-old (D) WT and AP-2 cKO mice (N=3 per genotype and age). Statistical significance was determined by two-way ANOVA followed by Tukeýs multiple comparison (2 months: lobule IV/V p<0.0001; lobule VI p<0.0001; lobule IX p=0.0406. 3 months: lobule III p<0.0001; lobule IV/V p<0.0001; lobule VI p; lobule VIII p<0.0001; lobule IX p=0.0011). **(E)** Schematic illustrating the experimental timeline for stereotactic injection of AAV1/2- Ef1α-DIO-EYFP into lobule IV/V of the cerebellum of AP-2 cKO or control littermates. **(F)** Representative confocal images of Purkinje cells from WT and AP-2 cKO mice expressing EYFP in a Cre-dependent manner (using AAV1/2-Ef1α-DIO-EYFP). Scale bar: 20µm, right insert 5µm, left insert 2µm. **(G)** Sholl diagram of dendritic arborisation within 10µm radius from the soma of EYFP-transduced WT and AP-2 cKO Purkinje cells (N=5 mice for WT, N=3 mice for KO). Statistical significance was determined by two-way ANOVA followed by Tukeýs multiple comparison. **(H)** Sholl-based analysis of total number of intersections in AP-2 cKO and WT EYFP-transduced Purkinje cells. Each dot represents one cell (WT: n=27 cells from N=5 mice; cKO: n=27 cells from N=3 mice). Statistical significance was determined by unpaired two-tailed Student’s t-test (WT: 803.4 ± 74.27, cKO: 560.9 ± 40.83; p=0.0061). **(I,J)** Quantification of the number of spines in proximal (I) and distal dendrites (J) of WT and AP-2 cKO EYFP-transduced Purkinje cells. Each dot represents one cell (n=18 cells, N=3 mice per genotype). Statistical significance was determined by unpaired two-tailed Student’s t-test (proximal dendrites WT: 19.87 ± 0.5185, cKO: 26.07 ± 0.7715; p<0.0001; distal dendrites WT: 19.96 ± 0.6461, cKO: 25.46 ± 0.6818; p<0.0001). Data is presented as mean±SEM. * p ≤0.05; ** p≤0.01; *** p≤0.001; **** p≤0.0001. n.s.-non-significant.

To take a detailed look at Purkinje cell morphology at the onset of their degeneration, we stereotactically delivered an adeno-associated virus (AAV) carrying EYFP flanked by double-inverted orientation (DIO) sites (AAV1/2-Ef1α-DIO-EYFP), allowing Cre-dependent expression, into the cerebellum of 4-week-old WT (*Ap2m1*^wt/wt^;L7*^Cre^*) and AP-2 cKO (*Ap2m1^fl/fl^*;L7*^Cre^*) mice (**Fig. 2E**). Analysis of 6-week-old EYFP-expressing Purkinje cells in AP-2 cKO mice revealed only slight changes in their branching pattern (**Fig. 2F-H**). Surprisingly, 6-week-old Purkinje cells lacking AP-2 showed a significant increase in spine-like protrusions emanating from both proximal and distal dendrites (**Fig. 2I,J**, see also Fig. 2F). The observed decrease in overall branching and the increase in spine density was likely independent of canonical function of AP-2 in CME, since this phenotype did not occur in cultured Purkinje cells treated with the clathrin inhibitor PitStop2 (**Fig. S3E-I**). These data provide the first line of evidence that synaptic dysfunctions may precede cell death and gait abnormalities in AP-2 cKO mice.

### AP-2-deficient Purkinje cells reveal proteome alterations at glutamatergic synapses

To further explore the precise role of AP-2 in Purkinje cells, we conducted quantitative mass spectrometry (MS) analysis of WT and AP-2 cKO cerebellar proteome before (1-month-old) and during (2-month-old) the “cell death” window. Only a few proteins were deregulated in the cerebellum of 1-month-old mice (**Fig. 3A**), whereas proteome changes were abundant at 2 months of age (**Fig. 3B, Table S1**). Notably, half of the deregulated proteins at 1 month overlapped with proteins that were altered in 2-month-old AP-2 cKO cerebellum (**Fig. S4A**). Next, we subjected proteins identified in the MS analysis at both ages to Gene Ontology (GO) term enrichment analysis aiming at identifying cellular components and biological processes, which are regulated by AP-2 in the cerebellum. Analysis of the upregulated proteome of the 1-month-old AP-2 cKO cerebellum revealed "glutamatergic synapse" as one of the highly ranked cellular GO components (**Fig. 3C**), suggesting that synaptic alternations may indeed precede Purkinje cell degeneration in our mouse model. Interestingly, in the 2-month-old cerebellum, most of the downregulated proteins were components of the “postsynapse” and “glutamatergic synapses” (**Fig. 3D**). Consistent with the onset of Purkinje cell loss at 2 months of age, but not at 1 month of age, “neuronal death” and “microglia activation” were the most prominently upregulated GO biological processes in the 2-month-old AP-2 cKO cerebellar proteome (**Fig. 3E**). The fact that apoptotic caspases 3 and 6 were upregulated in the 2-month-old AP-2 cKO proteome and in AP-2 cKO Purkinje cells (see Fig. 3B, **Fig. S4B,C**) suggests that AP-2 KO PCs undergo cell death by apoptosis. Moreover, in line with our behavioural data (see Fig. 1), we observed that many downregulated proteins in the cerebellum of 2-month-old AP-2 cKO mice are associated with SCA (**Fig. 3F**). Since these proteome alterations reflect “bulk” alterations in the cerebellum and can be attributed to changes in other neuronal types and glia, we conducted a series of experiments to specifically analyse the proteome of Purkinje cells lacking AP-2 using a combination of proximity labelling and affinity purification MS analysis. We stereotactically delivered an AAV carrying a cytosolic version of dimerized ascorbate peroxidase (dAPEX2) flanked by DIO (ssAAV-1/2-hEF1α-DIO-dAPEX2) into the 6-week-old cerebellum of WT (i.e. *Ap2m1*^wt/wt^;L7*^Cre^*) and AP-2 cKO (i.e. *Ap2m1^fl/fl^*; L7*^Cre^*) mice to allow Purkinje cell-dependent expression of APEX2 (**Fig. 3G**). In the presence of hydrogen peroxide, APEX biotinylates nearby proteins that can subsequently be enriched with streptavidin beads and identified by MS. Hydrogen peroxide treatment of acute cerebellar slices of mice sacrificed 3 weeks later after receiving a stereotactic injection of DIO-dAPEX2 AAV resulted in significant enrichment of streptavidin-labelled proteins (**Fig. S4D**). By analysing the streptavidin-enriched proteome in WT and AP-2 cKO cerebellum, we found a large number of proteins that were decreased in Purkinje cells lacking AP-2 compared to WT (**Fig. 3H, Table S2**). These downregulated proteins were mainly regulators of “actin filament organisation” and “synapse organisation” (**Fig. 3I**). In addition, several proteins with endocytosis-related functions were significantly decreased in AP-2 cKO Purkinje cells, including many direct players of the CME (DMN3, CLTC, SH3GL2, DNAKC6 and others) (**Fig. 3J**), confirming the robustness of our approach. By comparing the downregulated proteins in whole cerebellar proteome to the Purkinje-cell enriched proteome, we found that approximately one quarter of all downregulated proteins in the cerebellum of 2-month-old AP-2 cKO mice (see Fig. 3B,D,F) were specifically reduced in Purkinje cells (**Fig. 3K,L**). More importantly, these commonly downregulated proteins were associated with SCA, and majority were players at glutamatergic synapses (**Fig. 3L**). Taken together, our comprehensive proteomic analysis indicate that deletion of AP-2 in Purkinje cells leads to alterations at glutamatergic synapses.

**Figure 3.**
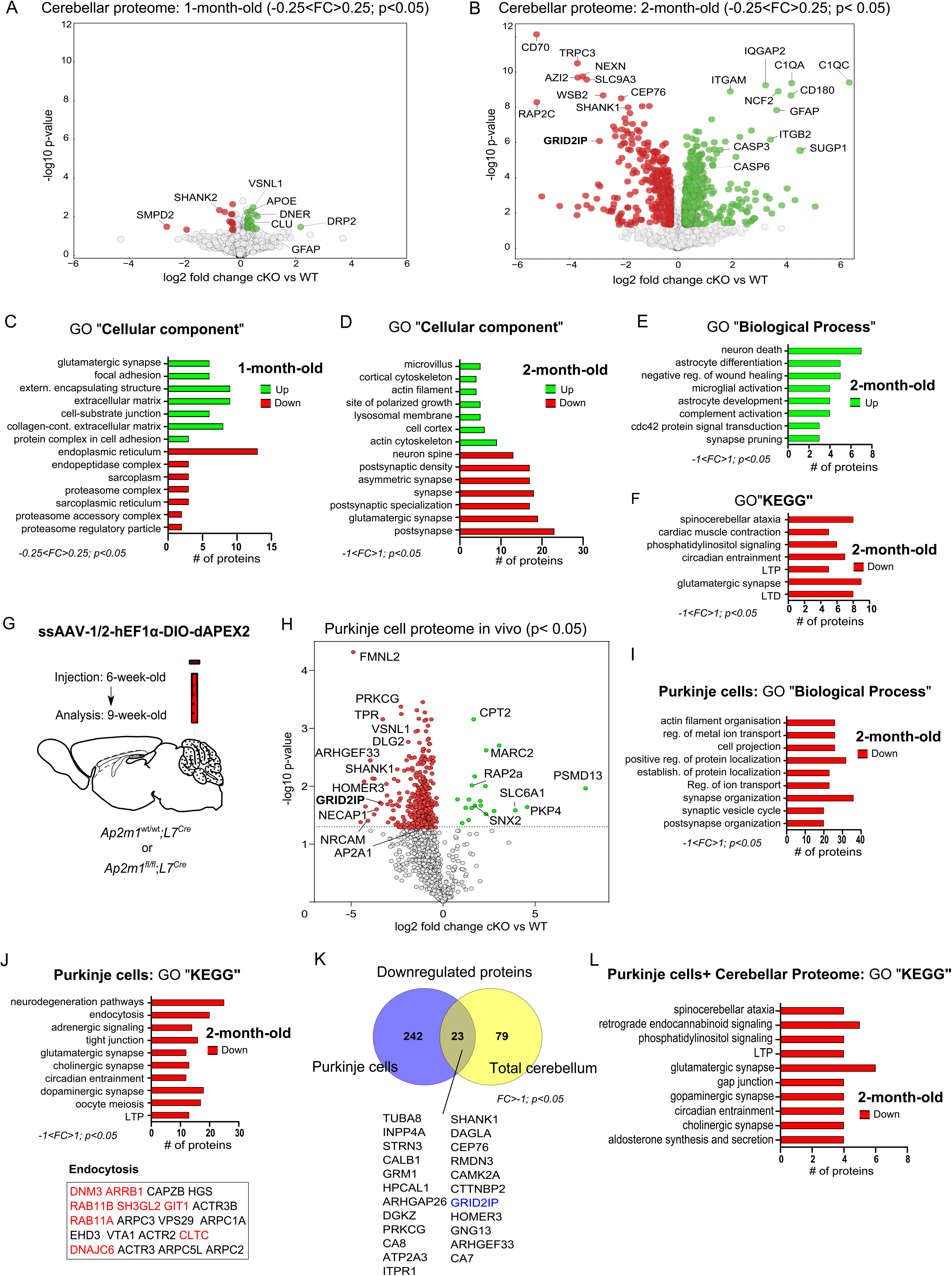
Proteomic analysis of AP-2 cKO cerebellum reveals early alterations at glutamatergic synapses. (**A,B**) Volcano plot of differentially expressed proteins in WT and AP-2 cKO cerebellar lysates at 1 month (A) and 2 months (B) of age analyzed using label-free proteomic approach. Red and green colored circles mark significantly upregulated and downregulated proteins at p < 0.05 and log2 fold change of >-0.25 and >0.25, respectively. See also Table S1. N=5 per genotype. **(C, D)** ShinyGo v0.80-based gene ontology (GO) analysis of deregulated “Cellular component”-enriched terms in the cerebellum of 1-month-old (C) and 2-month-old AP-2 cKO mice (D). **(E)** ShinyGo v0.80-based gene ontology (GO) analysis of upregulated “Biological process”-enriched terms in the cerebellum of 2-month-old AP-2 cKO mice. **(F)** ShinyGo v0.80-based gene ontology (GO) analysis of downregulated “KEGG”-enriched terms in the cerebellum of 2-month-old AP-2 cKO mice. **(G)** Schematic illustrating the experimental timeline for stereotactic injection of ssAAV-1/2- hEF1α-DIO-dAPEX2 into lobules IV/V of the cerebellum of AP-2 cKO or control littermates. **(H)** Volcano plot of differentially expressed proteins in dAPEX2-transduced Purkinje cells of WT vs AP-2 cKO mice. APEX2-biotinylated proteins were enriched by affinity pull-down and analyzed using label-free proteomic approach. Red and green colored circles mark significantly upregulated and downregulated proteins at p < 0.05, respectively. See also Table S2. N=2 per genotype. **(I)** ShinyGo v0.80-based gene ontology (GO) analysis of downregulated “Biological process”-enriched terms in 2-month-old AP-2 cKO Purkinje cells. **(J)** ShinyGo v0.80-based gene ontology (GO) analysis of downregulated “KEGG”-enriched terms in 2-month-old AP-2 cKO Purkinje cells. Lower panel indicates downregulated proteins within the term “Endocytosis”. Known binding partners of AP-2 are indicated in red. **(K)** Venn diagram of commonly downregulated proteins (p < 0.05 and log2 fold change of < > 1) in the total cerebellar proteome and APEX2-transduced 2-month-old AP-2 cKO Purkinje cells. **(L)** ShinyGo v0.80-based gene ontology (GO) analysis of commonly downregulated “KEGG”-enriched terms in 2-month-old total cerebellar proteome and APEX2-transduced AP-2 cKO Purkinje cells.

### AP-2 regulates the stability of the PF synapses-enriched protein GRID2IP

To reveal how the loss of AP-2 in Purkinje cells leads to early changes in synaptic organization, we performed a MS-based analysis of AP-2 interaction partners in the cerebellum using the AP-2α antibody as a bait (**Fig. 4A, Table S3**). Next to pulling-down components of the AP-2 complex and the known AP-2 binding partners (i.e. STON2, AAK1, CLTB, PICALM), we identified GRID2IP (also known as Delphilin) as a novel AP-2 binding partner in the cerebellum. GRID2IP was among the 15 proteins bound to AP-2α in our MS pull-down experiments and downregulated in AP-2 cKO Purkinje cells in the dAPEX2-based proteomic analysis (**Fig. 4B**). GRID2IP is a PDZ and formin homology (FH) domain-containing protein, selectively expressed in Purkinje cells (Miyagi et al., 2002). While FH domains are required to promote filamentous actin nucleation, PDZ domains are known to be present in proteins serving as scaffolds at postsynaptic membranes. In a series of biochemical and immunohistochemical experiments, we confirmed that AP-2 and GRID2IP interact and colocalize in Purkinje cells dendrites *in vivo* (**Fig. 4C-E**). Interestingly, mice with AP-2 deletion in Purkinje cells revealed significantly decreased levels of GRID2IP, both in cerebellar lysates (**Fig. 4F,G**, see also Fig.3B) and in Purkinje cells identified by calbindin immunostaining (**Fig. 4H**, see also Fig.3I). To map the distribution of GRID2IP on Purkinje cells in 3D, we reconstructed the dendritic trees of 6-week-old Purkinje cells from WT and AP-2 cKO mice injected with an AAV-EYFP-DIO at 4 weeks of age (see Fig. 2E). The density of GRID2IP distribution on Purkinje cell dendrites was significantly reduced in AP-2 cKO mice compared to controls (**Fig. 4I,J**). This loss of GRID2IP was not due to its reduced mRNA levels (**Fig. 4K,L**), suggesting that AP-2 might regulate the protein stability of GRID2IP in Purkinje cells. The ubiquitin-proteasome system is a central player in regulating turnover of cytoplasmic proteins, including the degradation of a homologous formin protein mDia2 (DeWard and Alberts, 2009). To test whether GRID2IP is more susceptible to proteasomal degradation in the absence of AP-2, we incubated WT and AP-2 cKO acute cerebellar slices with the proteasome inhibitor MG132. Indeed, a 6.5-hour treatment with MG132 was sufficient to increase GRID2IP levels in WT Purkinje cells and restore its normal expression in Purkinje cells lacking AP-2 (**Fig. 4M,N**), suggesting that the binding of AP-2 to GRID2IP might be required to protect GRID2IP from proteasome degradation.

**Figure 4.**
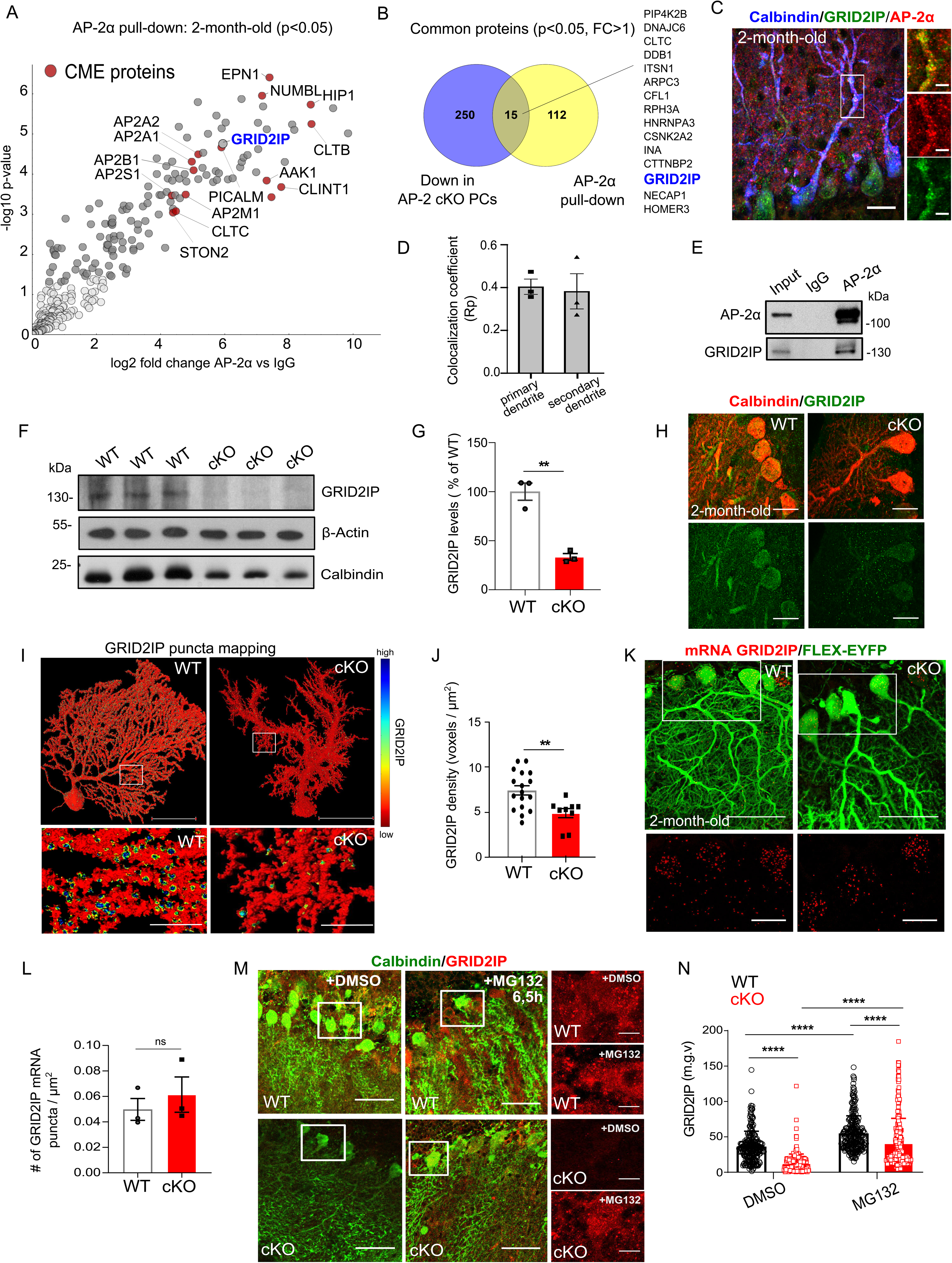
GRID2IP is a novel binding partner of AP-2 in the cerebellum. **(A)** Volcano plot showing interaction partners identified by pull-down of AP-2α from the cerebellum of 2-month-old WT mice (N=5) followed by quantitative mass spectrometry. Grey circles indicate significantly enriched proteins in the AP-2α pull-down compared to the IgG control. Known AP-2 interaction partners with a function in CME are highlighted in red. See also Table S3. **(B)** Venn diagram of proteins significantly downregulated in AP-2 cKO PCs (p < 0.05 and log2 fold change of > 1) and found to be significantly enriched in AP-2α pull-down in WT cerebellum (p < 0.05 and log2 fold change of > 1). **(C)** Representative confocal image of cerebellar Purkinje cells from 2-month-old WT mice stained with antibodies against AP-2α (red), GRID2IP (green) and Calbindin (blue). Scale bar: 20µm, insert 5µm. **(D)** Colocalization analysis (Pearsońs coefficient) of AP-2α and GRID2IP in primary and secondary dendrites of Purkinje cells from 2-month-old WT mice (N=3). **(E)** Co-immunoprecipitation of endogenous AP-2α with GRID2IP in 2-month-old WT cerebellum lysates (N=3). Input 1,5% of the lysate. **(F,G)** Immunoblot analysis of GRID2IP levels in the cerebellum of 2-month-old WT and cKO mice. β-actin and Calbindin were used both as a loading control and for normalisation of protein levels. GRID2IP protein levels are significantly reduced in the cerebellum of 2-month-old AP-2 cKO mice compared to WT control set to 1. Each dot represents one mouse (N=3 per genotype). Statistical significance was determined by one-tailed unpaired Student’s t-test (cKO: 0.33.45 ± 3.44; p=0.0011). **(H)** Representative confocal images of Purkinje cells from 2-month-old WT and AP-2 cKO mice stained with antibodies against Calbindin (red) and GRID2IP (green). Scale bar: 20µm. **(I)** AMIRA-based visualisation of GRID2IP distribution in 6-week-old Purkinje cells of WT and cKO mice. 3D reconstruction of Purkinje cells is based on GFP immunostaining after transduction with AAV1/2-Ef1α-DIO-EYFP (see Fig. 2E). Scale bar, 50µm, insert 10µm. **(J)** Quantification of GRID2IP voxels in Purkinje cells dendrites reveals a decrease in GRID2IP protein expression in AP-2 cKO mice compared to WT. Each dot represents one cell (WT: n=17 cells from N=3 mice; cKO: n=9 cells from N=3 mice). Statistical significance was determined by unpaired two-tailed Student’s t-test (WT: 7.5 ± 0.52; cKO: 4.9 ± 0.51; p=0.0043). **(K)** Immunofluorescent staining for GFP and multiplex fluorescent *in situ* hybridization for *Grid2ip* mRNA on cerebellar sagittal sections of 6-week-old WT and cKO mice transduced with AAV1/2-Ef1α-DIO-EYFP. Scale bar, 50µm, insert, 20µm. **(L)** Quantification of *Grid2ip* mRNA puncta reveals no significant difference in Purkinje cells of WT and AP-2 cKO mice. Each dot represents one mouse (WT: 0.050±0.008, sKO: 0.061±0.014, p=0.514, N=3 mice per genotype). Statistical significance was determined by unpaired two-tailed Student’s t-test. **(M)** Representative confocal images of acute cerebellar slices from 2-month-old WT and AP-2 cKO mice stained with antibodies against Calbindin (green) and GRID2IP (red). Acute slices were treated with 100µM MG132 for 6.5h. Equal amount of DMSO was used as a control. Scale bar: 50µm, inserts 10µm. **(N)** Quantification of GRID2IP protein levels in WT and AP-2 cKO cerebellar acute slices treated for 6.5h with either 100 µM of MG132 or equal amount of DMSO. Each dot represents one cell (WT DMSO: n= 233 cells from N=4 mice; WT MG132: n= 253 cells from N= 4 mice; cKO DMSO: n= 233 cells from N=4 mice; cKO MG132: n= 249 cells from N=4 mice). Statistical significance was determined by two-way ANOVA followed by Tukeýs multiple comparison test (WT DMSO: 36.17 ± 1.424, WT MG132 54.86 ± 1.566; p<0.0001. cKO DMSO: 13.02 ± 0.8310, cKO MG132: 41.28 ± 2.208; p<0.0001 for comparison with WT DMSO and cKO MG132). Data is presented as mean±SEM. * p ≤0.05; ** p≤0.01; *** p≤0.001; **** p≤0.0001. n.s.-non-significant.

### Loss of AP-2 leads to synaptic accumulation of GLUR**δ**2 and causes an imbalance of PF and CF synapses

Our data above describe a previously unknown interaction between AP-2 and the postsynaptic scaffolding protein GRID2IP (**Fig. 5A**). Since GRID2IP was originally discovered as a glutamate δ2 receptor (GLURδ2, also known as GluD2, encoded by *GRID2* gene)-interacting protein (Miyagi et al., 2002), an orphan glutamate receptor selectively expressed at PF-Purkinje cell synapses (Lomeli et al., 1993; Takayama et al., 1995), we first analyzed the levels of GLURδ2 at WT and AP-2 cKO PF synapses identified by their VGLUT1 expression (Fremeau et al., 2001). We found that GLURδ2 levels were upregulated in the dendrites of 4-week-old AP-2 cKO Purkinje cells (a period that precedes Purkinje cell loss) (**Fig. 5B,C**), a phenotype particularly evident at the PF synapses (**Fig. 5D**) and persistent until 6 weeks of age (**Fig. S5A**). Although GLURδ2 is structurally similar to ionotropic glutamate receptors, it is not gated by glutamate and no channel activity has yet been identified. Nevertheless, mice lacking GLURδ2 show impaired long-term depression (LTD) and motor learning, while a homozygous *GRID2* mutation in humans results in SCA18 (Hills et al., 2013; Kashiwabuchi et al., 1995; Panda et al., 2022). These motor deficits are explained by the synaptogenic function GLURδ2 at PFs, where it acts as a receptor for cerebellin-1 (Cbln1) secreted by granule cells (Matsuda et al., 2010; Uemura et al., 2010). Cbln1, in turn, binds neurexin, a presynaptic cell adhesion protein on PFs (Yuzaki, 2017) (see Fig. 5A). In support of its synaptogenic role, GLURδ2 KO mice lose nearly half of all PF synapses (Hideo et al., 1997; Ichikawa et al., 2002). Conversely, GLURδ2 overexpression leads to an excess of PFs and numerous spine-like protrusions (Takeo et al., 2021). Taken into account the accumulation of GLURδ2 at AP-2 cKO PF synapses, we speculated that this accumulation leads to increased PF formation. We analyzed PF synapses on 6-week-old WT and AP-2 cKO Purkinje cells by performing 3D analysis of VGLUT1 distribution on reconstructed Purkinje cell dendrites. Indeed, we observed that deletion of AP-2 increased PF synapse density (**Fig. 5E,F,I**). The increase in VGLUT1-positive synapses in Purkinje cells lacking AP-2 was evident as early as 4 weeks of age, consistent with our data described above (see Fig. 3C), providing further evidence that dysfunction at glutamatergic synapses precedes the loss of Purkinje cells and gait abnormalities in our model. Strikingly, these changes were accompanied by a nearly complete loss of CF synapses on Purkinje cell dendrites (**Fig. 5G,H,J**) and an accumulation of perisomatic CF synapses in AP-2 cKO mice (**Fig. 5K,L**). Taken together, our data suggest that AP-2 localized in Purkinje cells is required for the refinement of PF/CF synapses in the cerebellum.

**Figure 5.**
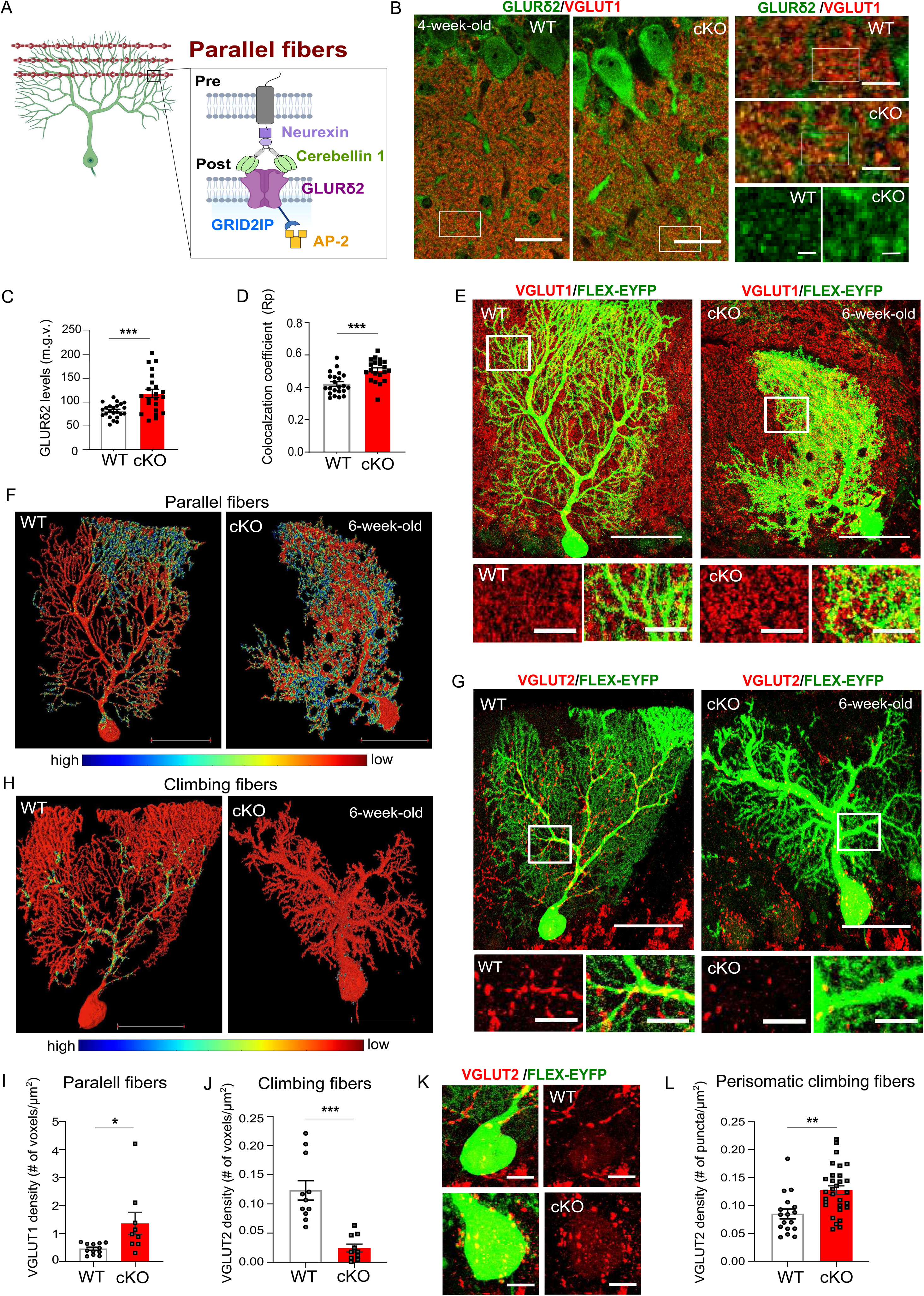
Reorganization of PF and CF inputs in AP-2 cKO cerebellum. **(A)** Schematic illustrating the interplay between GLURδ2, GRID2IP and AP-2 at Purkinje cell-PF synapses. **(B)** Representative confocal images of 1-month-old WT and AP-2 cKO cerebellum stained with antibodies against VGLUT1 (red) to mark PF synapses and GLURδ2 (green). Scale bar: 50µm, inserts upper panel 10µm, inserts lower panel 2µm. **(C)** Analysis of GLURδ2 protein levels in distal Purkinje cells dendrites reveals a significant increase in GLURδ2 levels in AP-2 cKO compared to WT. Each dot represents one image (WT: n=23 images from N=3 mice, cKO n=22 images from N=3 mice). Statistical significance was determined by unpaired two-tailed Student’s t-test (WT: 81.91 ± 3.121, cKO: 118.5 ± 8.751; P=0.0002). **(D)** Colocalization coefficient (Pearsońs correlation) analysis reveals a significantly increased colocalization between GLURδ2 and VGLUT1 at PF synapses of AP-2 cKO Purkinje cells compared to WT. Each dot represents one image (WT: n=22 images from N=3 mice, cKO: n=20 images from N=3 mice). Statistical significance was determined by unpaired two-tailed Student’s t-test (WT: 0.4202 ± 0.01424, cKO: 0.5044 ± 0.01561; p= 0.0003). **(E)** Immunofluorescence staining for GFP and the PF marker VGLUT1 (red) on sagittal cerebellar sections from 6-week-old WT and AP-2 cKO mice injected with AAV1/2-Ef1α-DIO-EYFP (see Fig. 2E). Scale bar: 50µm, inserts 10µm. **(F)** AMIRA-based 3D visualization of VGLUT1 distribution in 6-week-old Purkinje cells of WT and cKO mice. 3D reconstruction of Purkinje cells is based on GFP immunostaining after transduction with AAV1/2-Ef1α-DIO-EYFP. Scale bar: 50µm. **(G)** Immunofluorescence staining for GFP and the CF marker VGLUT2 (red) on sagittal cerebellar sections from WT and cKO mice injected with AAV1/2-Ef1α-DIO-EYFP (see Fig. 2E). Scale bar: 50µm, inserts 10µm. **(H)** AMIRA-based 3D visualization of VGLUT2 distribution in 6-week-old Purkinje cells of WT and cKO mice. 3D reconstruction of Purkinje cells is based on GFP immunostaining after transduction with AAV1/2-Ef1α-DIO-EYFP. Scale bar: 50µm. **(I)** Quantification of VGLUT1 voxels onto Purkinje cells dendrites reveals a significant increase in VGLUT1 puncta on AP-2 cKO Purkinje cell dendrites compared to WT. Each dot represents one cell (WT: n=11 cells from N=4 mice; cKO: n=9 cells from N=3 mice). Statistical significance was determined by unpaired two-tailed Student’s t-test (WT: 0.46 ± 0.056, cKO: 1.4 ± 0.40; p=0.0169). **(J)** Quantification of VGLUT2 voxels onto Purkinje cells dendrites reveals a significant decrease in VGLUT2 puncta on AP-2 cKO Purkinje cell dendrites compared to WT. Each dot represents one cell (WT: n=11 cells from N=3 mice; cKO: n=10 cells from N=3 mice). Statistical significance determined by unpaired two-tailed Student’s t-test (WT: 0.12 ± 0.017, cKO: 0.024 ± 0.0067; p<0.0001). **(K)** Cell bodies of WT and AP-2 cKO Purkinje cells magnified from Fig. 5G stained with an antibody against VGLUT2 (red) to mark CF synapses. Scale bar: 5µm. **(L)** Quantification of perisomatic VGLUT2 puncta in 6-week-old WT and AP-2 cKO Purkinje cells. Each dot represents one cell (WT: n=17 cells from N=3 mice; cKO: n=30 cells from N=3 mice). Statistical significance determined by unpaired two-tailed Student’s t-test (WT: 0.085 ± 0.009, cKO: 0.127 ± 0.008; p=0.0012). Data is presented as mean±SEM. * p ≤0.05; ** p≤0.01; *** p≤0.001; **** p≤0.0001.

Importantly, this regulation of PF/CF inputs by the AP-2 is achieved Purkinje cell-autonomously.

### AP-2 balances PF and CF synapses during development, but not in the adult brain

At birth, the soma of an individual Purkinje cell is innervated by multiple CFs (Hashimoto and Kano, 2013), whereas adult Purkinje cells receive input from one or a few CFs (Busch and Hansel, 2023). An activity-dependent regression of supernumerary CFs happens shortly after birth and is completed by postnatal day 21 (Jennifer et al., 2013). During this time period, the site of CF innervation undergoes a change, shifting from Purkinje cell soma to dendrites, a phenomenon termed as “CF translocation” (Hashimoto et al., 2009; Scelfo et al., 2003). Our data described above are consistent with the hypothesis that AP-2 regulates PF formation and that its loss results in overabundance of PF synapses, which interferes with CF translocation during early postnatal cerebellar development (**Fig. 6A**). If this hypothesis is correct, deletion of AP-2 after postnatal day 21 should not prevent CF translocation because their territory is already fully established. To test this hypothesis, we first analyzed the density of CFs at the Purkinje cell soma from 4-week-old WT and AP-2 KO mice. We found that AP-2 deletion during early postnatal development (as *L7-Cre* recombinase is fully active as early as 2 weeks after birth) resulted in a large increase in the number of perisomatic CFs, a phenotype consistent with their failure to translocate to the Purkinje cell dendrites (**Fig. 6B,C**). Next, we tested whether acute deletion of AP-2 in adult Purkinje cells also alters CF density. We stereotactically injected either an AAV expressing *L7* promoter-driven EYFP and Cre-recombinase (AAV2/rh10-L7-6-EGFP-P2A-Cre-WPRE) or a control AAV expressing EYFP under the *L7* promoter (AAV2/rh10-L7-6-EGFP-WPRE) in the cerebellum of 6-week-old of *Ap2m1^flox/flox^*mice (**Fig. 6D**). Analysis of 10-week-old EYFP-expressing AP-2-deficient Purkinje cells (**Fig. S5D,E**) showed no changes in their overall morphology (**Fig. 6E**) and/or in the number of their dendritic spines compared to controls (**Fig. 6F,G)**. Importantly, we observed that the CF territory was unaltered in Purkinje cells with acute deletion of AP-2 in adulthood (**Fig. 6H,I**), consistent with our hypothesis that AP-2 balances PF and CF synapses during early postnatal cerebellar development.

**Figure 6.**
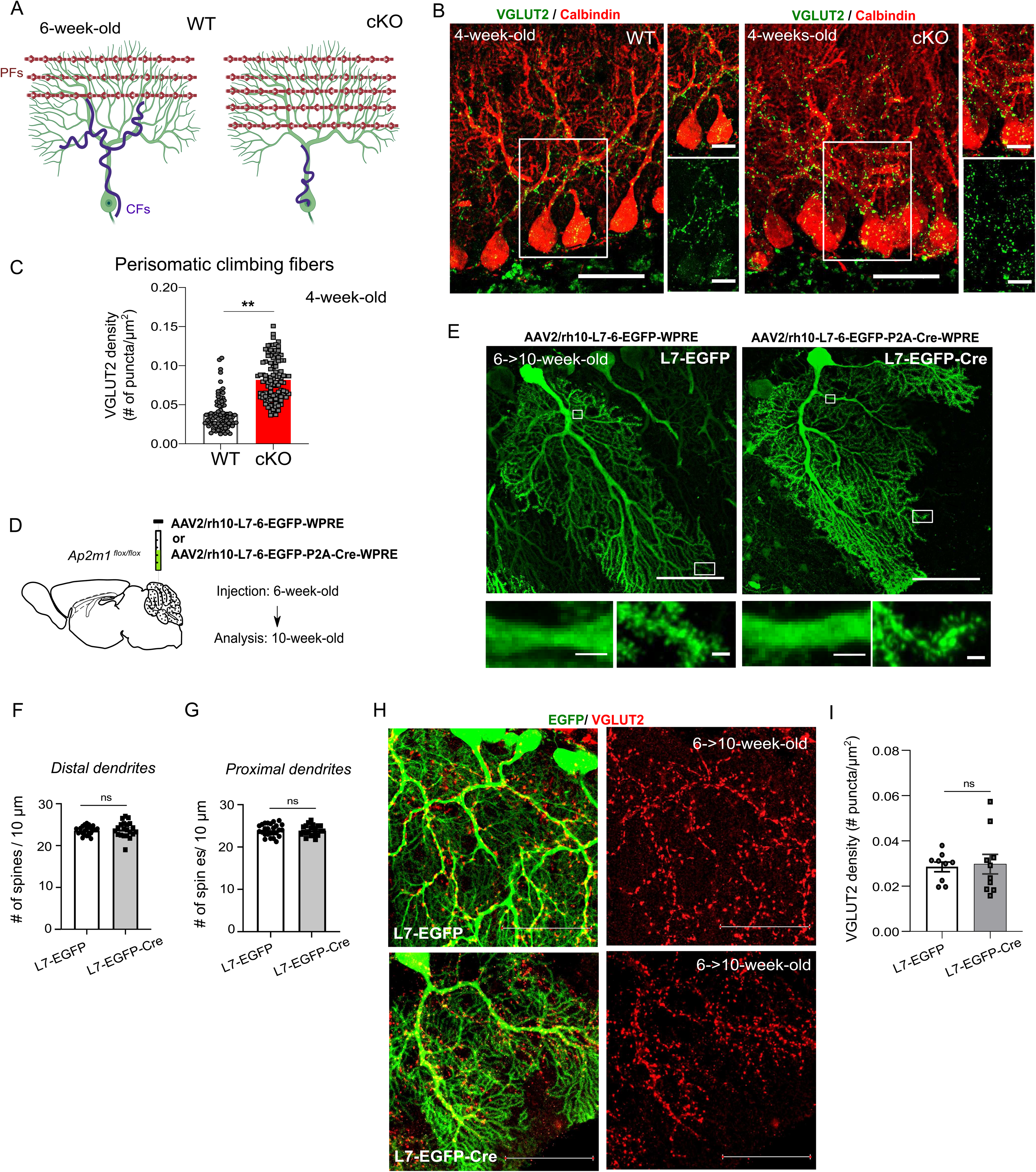
Rewiring of CFs is already present in the 4-week-old AP-2 cKO cerebellum but absent upon acute deletion of AP-2µ in adult mice. **(A)** Schematic illustrating the reorganisation of PF (brown) and CF (magenta) synapses in AP-2 cKO cerebellum. **(B)** Representative confocal images of 1-month-old WT and AP-2 cKO cerebellum immunostained for a CF marker VGLUT2 and Calbindin. Scale bar: 50µm, inserts 10µm. **(C)** Quantification of perisomatic VGLUT2 puncta in 4-week-old WT and AP-2 cKO Purkinje cells. Each dot represents one cell (WT: n=17 cells from N=3 mice; cKO: n=30 cells from N=3 mice). Statistical significance determined by unpaired two-tailed Student’s t-test (WT: 0.085 ± 0.009, cKO: 0.127 ± 0.008; p=0.0012). **(D)** Schematic illustrating the experimental timeline adopted for acute deletion of AP-2µ in the cerebellum by stereotactic injection of AAV2/rh10-L7-6-EGFP-P2A-Cre-WPRE (L7-EGFP-Cre) or control virus AAV2/rh10-L7-6-EGFP-WPRE (L7-EGFP) in the cerebellum of 6 week-old *Ap2m1^flox/flox^* animals. **(E)** Representative confocal images of 10-week-old Purkinje cells transduced with either L7-EGFP-Cre (resulting in acute AP-2 cKO) or control virus L7-EGFP. Scale bar: 50µm, inserts 10µm and 2µm. **(F,G)** Quantification of number of spines in Purkinje cells from *Ap2m1^flox/flox^* mice injected with either L7-EGFP control virus or L7-EGFP-Cre shows no difference in spine numbers after AP-2µ acute deletion in both proximal (F) and distal dendrites (G). Each dot represents one cell (Proximal dendrites, L7-EGFP: 24 cells from N=3 mice; L7-EGFP-Cre: n=21 cells from N=3 mice. Distal dendrites, L7-EGFP: 26 cells from N=3 mice; L7-EGFP-Cre: n=21 cells from N=3 mice). Statistical significance determined by unpaired two-tailed Student’s t-test. **(H,I)** No changes were detected in climbing fibres input after AP-2µ acute deletion, as revealed by VGLUT2 immunofluorescent staining (H) and analysis of its puncta density (I) (L7-EGFP: 0.028±0.002, L7-EGFP-Cre: 0.030±0.004, p=0.811, n=9 images from L7-EGFP and n=10 images from L7-EGFP-Cre-injected mice (each with >5 cells), N=3 mice for each conditions). Statistical significance determined by unpaired two-tailed Student’s t-test. Data is presented as mean±SEM. * p ≤0.05; ** p≤0.01; *** p≤0.001; **** p≤0.0001. n.s.-non-significant.

### Activity-dependent potentiation of Purkinje cell network activity in the absence of AP-2

Finally, we investigated whether an imbalance of PF/CF inputs in AP-2 mice alters Purkinje cell network activity. First, we evaluated the intrinsic electrophysiological properties of Purkinje cells in acute cerebellar slices. We found that Purkinje cells from 4-week-old AP-2 cKO mice had unaltered baseline intrinsic properties (**Fig. S6A-C**), but were less excitable compared to Purkinje cells from WT mice as they required a significantly higher intracellular current injection to elicit action potentials (**Fig. 7A-C**). In a second step, we investigated the output of WT and AP-2 cKO Purkinje cells in response to synaptic activity. During motor learning, Purkinje cells fire action potentials at frequencies around 100 Hz and higher (Gilbert and Thach, 1977). To study the synaptic response of Purkinje cells to a 100 Hz stimulation in WT and AP-2 cKO mice, we employed GCaMP7f-based live imaging in organotypic cerebellar slices, prepared at postnatal day 8 and imaged at day-in-vitro 21, before Purkinje cells undergo degeneration. Surprisingly, we observed that both the average response to stimulation and the peak amplitude of responses were higher in Purkinje cells from AP-2 cKO animals than in Purkinje cells from WT mice (**Fig. 7D-G, Fig. S6D**). Moreover, Ca^2+^ transients of AP-2 cKO Purkinje cells were not only significantly increased within 1-10 sec post-stimulation (**Fig. 7H**), but also remained significantly higher than in WT cells up to 20 sec after stimulation at 100Hz (**Fig. 7I**). In line with this, the average time to reach the maximal peak response was delayed in AP-2 cKO Purkinje cells (**Fig. 7J, Fig. S6E**), likely as a result of the second, lower amplitude peak of Ca^2+^ transients present in these cells (see Fig. 7G). These results may indicate that more recurrent network activity propagates through the cerebellar network in AP-2 cKO mice in the period before Pukinje cell degeneration. At the same time, the stereotypicity of stimulus-evoked neuronal activity was significantly lower in Purkinje cells from cKO mice compared to the WT (**Fig. 7K,L**). Taken together, these data suggest that deletion of AP-2 leads to asynchronous firing and activity-dependent potentiation of Pukinje cell activity.

**Figure 7.**
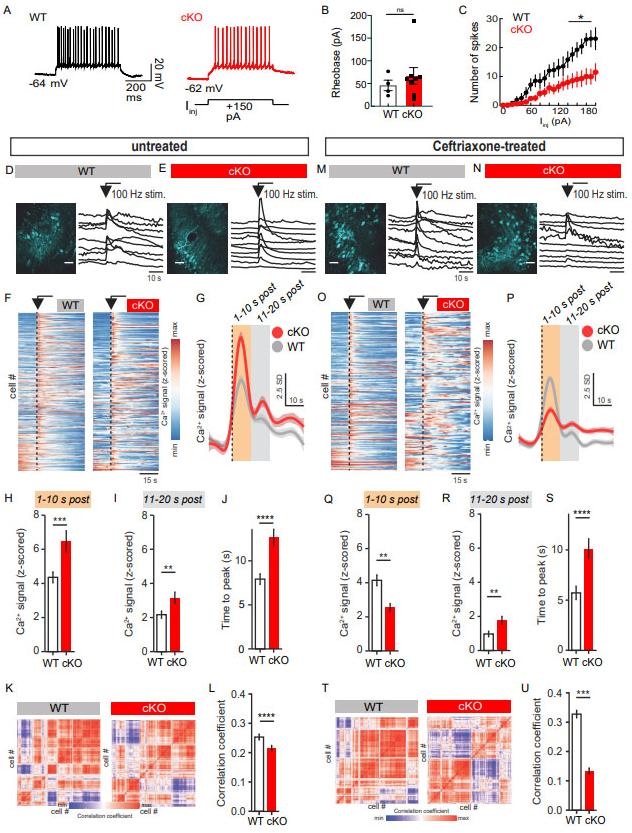
AP-2 functions in Purkinje cells to regulate cerebellar network excitability. **(A)** Example whole-cell recordings of Purkinje cells in acute cerebellum slices in response to 150 pA of intracellular injection current injection from 4-week-old WT and AP-2 cKO mice. **(B)** Analysis of Purkinje cell intrinsic properties in 4-week-old WT and AP-2 cKO mice. No significant changes were detected in cKO Purkinje cell rheobase (WT n=4 cells from N=3 mice, cKO n=8 cells from N=2 mice). Statistical significance was determined by two-tailed Mann-Whitney test. The raw data of the patch-clamp recordings are listed in Table S5. **(C)** Lower number of spikes evoked by intracellular current injection in Purkinje cells of cKO (n=8 neurons; N=2 mice) compared to WT animals (n=4 neurons from N=3 mice). Statistical significance was determined by Mann-Whitney two-tailed test. The raw data of the patch-clamp recordings are listed in Table S5. **(D,E)** Representative responses of Purkinje cells transfected with ssAAV-9/2-mCaMKIIα-jGCaMP7f-WPRE-bGHp(A) in WT (D) and AP-2 cKO (E) cerebellar organotypic slices to 100 Hz stimulation for 1 sec (i.e. 100 action potentials). Each trace is a representative of one cell within field of view. Scale bar, 50µm. **(F)** Activity map across all neurons sorted by maximum response to stimulation. **(G-I)** Peri-stimulus response (G) and quantification of response during either 1-10s (H) or 11-20s (I) post-stimulation with 100 Hz of Purkinje cells derived from WT or AP-2 cKO cells cerebellar organotypic slices (H, WT: 4.356±0.204, cKO: 6.469±0.405, p=0.00023, I, WT: 4.138±0.249, cKO: 2.544±0.199, p=0.0013). Data are from N=6 WT and N=5 cKO mice, with 457 and 255 Purkinje cells, accordingly. Statistical significance was determined by Welch’s t-test. **(J)** Analysis of the time required to reach the maximal Ca2+ response elicited by 100Hz stimulation in WT and AP-2 cKO Purkinje cells transduced with ssAAV-9/2-mCaMKIIα-jGCaMP7f-WPRE-bGHp(A) (WT: 7.926±0.376, cKO±12.641±0.798, p<0.0001). See Fig. S6B for the illustration of the point used for the analysis in WT and AP-2 cKO cells. Data are from N=6 WT and N=5 cKO mice, with 445 and 251 Purkinje cells, accordingly. Statistical significance was determined by Welch’s t-test. **(K,L)** Correlation matrix (K) and quantification (L) of stimulus-evoked responses across all WT (left) and AP-2 cKO Purkinje cells (right) (WT: 0.253±0.012, KO: 0.215±0.013, p<0.0001). Data are from N=6 WT and N=5 cKO mice, with 457 and 255 Purkinje cells, accordingly. Statistical significance was determined by Welch’s t-test. **(M,N)** Representative responses of Purkinje cells transfected with AAV-CamKIIα-GCaMP7f in WT (M) and AP-2 cKO (N) cerebellar organotypic slices treated with 100 µM ceftriaxone for 7 days starting at day in vitro (DIV) 15 and subjected to 100 Hz stimulation for 1sec (i.e. 100 action potentials). Each trace is a representative of one cell within field of view. Scale bar, 50µm. **(O)** Activity map across all neurons treated with 100 µM ceftriaxone, sorted by maximum response to stimulation. **(P-R)** Peri-stimulus response (P) and quantification of response during either 1-10s (Q) or 11-20s (R) post-stimulation with 100Hz of Purkinje cells derived from WT or AP-2 cKO cells cerebellar organotypic slices treated for 7 days with 100 µM ceftriaxone (Q, WT: 4.138±0.2249, cKO: 2.544±0.199, p=0.0015, R, WT: 0.971±0.058, cKO: 1.762±0.138, p=0.002). Data are from N=4 WT and N=2 cKO mice, with 277 and 164 Purkinje cells, accordingly. Statistical significance was determined by Welch’s t-test. **(S**) Analysis of the time required to reach the maximal Ca2+ response elicited by 100Hz stimulation in WT and AP-2 cKO Purkinje cells transfected with ssAAV-9/2-mCaMKIIα-jGCaMP7f-WPRE-bGHp(A) and treated with 100 µM ceftriaxone (WT: 5.721±0.353, cKO±10.116±0.813, p<0.0001). Data are from N=4 WT and N=2 cKO mice, with 262 and 155 Purkinje cells, accordingly. Statistical significance was determined by Welch’s t-test. **(T,U)** Correlation matrix (T) and quantification (U) of stimulus-evoked responses across all WT (left) and AP-2 cKO cells (right) after 100µM ceftriaxone treatment. (WT: 0.328±0.020, KO: 0.134±0.010, p<0.0001). Data are from N=4 WT and N=2 cKO mice, with 277 and 164 Purkinje cells, accordingly. Statistical significance was determined by Welch’s t-test. Data is presented as mean±SEM. * p ≤0.05; ** p≤0.01; *** p≤0.001; **** p≤0.0001. n.s.-non-significant.

To test whether excess synaptic input due to increased PF abundance (see Fig. 5I) is the cause of AP-2 cKO Purkinje cell hyperexcitation, we treated organotypic slices with ceftriaxone, a β-lactam antibiotic known to facilitate synaptic glutamate reuptake via its transcriptional upregulation of the glial glutamate transporter GLT-1 (Rothstein et al., 2005) (encoded by the *Slc1a2* gene, also known as EAAT2) (**Fig. 7M-O**). GLT-1 is expressed in Bergmann glia (Rothstein et al., 1994) and is responsible for more than 90% of total glutamate uptake in the brain (Kim et al., 2011). The transcriptional upregulation of GLT-1 via ceftriaxone is known to provide neuroprotection in various animal models of excitotoxicity (Leung et al., 2012; Rebec, 2013; Rothstein et al., 2005). Ceftriaxone treatment of organotypic slice culture for 7 days was sufficient to increase GLT-1 levels in the cerebellum (**Fig. S6F,G**) and markedly reduce network excitation in AP-2 cKO Purkinje cells (**Fig. 7P,Q, Fig. S6H)**. Of note, GLT1 levels were not altered *per se* in cerebellar lysates from AP-2 cKO mice (**Fig. S6 I,J**). Interestingly, the time to peak response continued to be delayed in AP-2 cKO Purkinje cells treated with ceftriaxone (**Fig. 7S**), although it improved slightly compared to the untreated condition (see Fig. 7J). This delay is likely due to an increase in the number of perisomatic CF synapses in AP-2 cKO mice (see Fig. 6L). Additionally, ceftriaxone treatment further decreased the stereotypicity of stimulus-evoked responses in cKO cells, resulting in a significant, almost threefold reduction in the cKO group compared with the WT group (**Fig. 7T,U**). Overall, our data suggest that early PF/CF input rewiring in AP-2 cKO Purkinje cells correlates with their hyperexcitability and that facilitation of synaptic glutamate reuptake normalizes their network excitation. On the other hand, the exacerbation of asynchronous firing of AP-2 cKO Purkinje cells after ceftriaxone treatment (see Fig.7U) is likely due to the loss of dendritic CFs in these mice, consistent with the previously described role of CF synapses in inhibiting neighbouring Purkinje cells via ephaptic coupling (Han et al., 2020).

## Discussion

Our findings provide significant insights into the critical role played by the endocytic adaptor AP-2 in shaping cerebellar circuitry. Through a series of experiments utilizing a novel mouse model with Purkinje cell-specific deletion of AP-2, we demonstrate that AP-2 plays a pivotal role in maintaining the delicate balance between PF and CF synapses, which is essential for Purkinje cell function. The absence of AP-2 leads to Purkinje cell hyperexcitation, subsequent neurodegeneration, and severe impairment of motor coordination. AP-2 is well-known for its involvement in CME (Conner and Schmid, 2003), a fundamental process that in neurons regulates synaptic vesicle recycling and neurotransmitter receptor trafficking (Camblor-Perujo and Kononenko, 2022; Kononenko and Haucke, 2015). While several studies have investigated the role of AP-2 in dendritogenesis and cargo sorting in various neuronal populations (Garafalo et al., 2015; Kononenko et al., 2017; Koscielny et al., 2018; Li et al., 2016), its role in synaptogenesis and neuronal circuitry formation remains largely unexplored. Our findings represent the first elucidation of AP-2 importance in orchestrating synaptic connectivity, suggesting its previously unrecognized role in shaping neuronal circuits.

The observed phenotypic changes in AP-2 cKO Purkinje cells suggest a cerebellum-specific function. While the reduction in dendritic complexity aligns with previous findings in cortical and hippocampal neurons (Kononenko et al., 2017; Koscielny et al., 2018), the fully established dendritic trees and increased spinogenesis of 2-month-old AP-2 cKO Purkinje cells suggest that AP-2 in the cerebellum may not be directly involved in the initial development of dendrite morphology, but instead exert regulatory control over synapse formation during early Purkinje cell maturation. This cerebellum-specific function of AP-2 is likely due to the unique architecture of the cerebellum, where precise regulation of synaptic connectivity is established postnatally (Leto et al., 2016) and can be mechanistically explained by its interaction with GRID2IP, a postsynaptic scaffold selectively enriched at PF-Purkinje cell synapses. GRID2IP functions in Purkinje cells by linking the GLURδ2 receptor, a master of PF synaptogenesis (Hideo et al., 1997; Ichikawa et al., 2002; Matsuda et al., 2010; Uemura et al., 2010), to the actin cytoskeleton. Mutations in GLURδ2 are associated with cerebellar ataxia in patients (Panda et al., 2022) and mice, including *lurcher* and *hotfoot* mutants (Miyoshi et al., 2014; Zuo et al., 1997). Both CF and PF fibers are severely altered in GLURδ2 KO mice, as the distal PC dendrites are innervated by CFs instead of PFs (Hideo et al., 1997; Ichikawa et al., 2002; Kashiwabuchi et al., 1995). We have shown that the intricate interplay between AP-2 and GRID2IP prevents proteasomal degradation of GRID2IP and that loss of AP-2 leads to accumulation of GLURδ2 at PF synapses, promoting their synaptogenesis. The exact mechanism by which GRID2IP regulates GLURδ2 localization at PF synapses remains to be determined. It is possible that the mechanisms are related to its role in lateral trafficking of GLURδ2, similar to what has been described for β-III spectrin-dependent regulation of EAAT4 trafficking (Ikeda et al., 2006). Nonetheless, our finding on AP-2 as a negative regulator of cerebellar PF synaptogenesis challenges the conventional view of AP-2 role solely in CME and synaptic vesicle recycling and suggests a more complex involvement in regulating synaptic development and plasticity in the brain.

One of the key observations of this study is the severe and early ataxia (first appearing around 7 weeks of age) exhibited by mice lacking AP-2 in Purkinje cells. This phenotype is associated with aberrant synaptic rewiring characterized by increased PF synapses and decreased CF inputs. Our data suggest that synaptic alteration precede Purkinje cell loss and that dysregulation of synaptic connectivity might contribute to motor coordination problems in our mouse model. We propose that elevated PF synaptogenesis is the primary event causing network hyperexcitation and possibly Purkinje cell degeneration as a result of glutamate excitotoxicity, similar to what has been described in SCA mouse models and induced pluripotent stem cells from SCA patients (Chuang et al., 2019; Custer et al., 2006). Several lines of evidence support our hypothesis that enhanced synaptogenesis of PF synapses is a key event in the cascade of neurodegeneration occurring in AP-2 cKO mice. First, AP-2 cKO cerebella exhibit increased PF innervation of distal dendrites as early as 4 weeks of age, accompanied by increased enrichment of GLURδ2 at PF synapses and subsequent increased spinogenesis in 6-week-old AP-2 cKO Purkinje cells. Second, the number of CF synapses at the somata of 4-week-old AP-2 cKO Purkinje cells is high, and beyond that point, CF terminals fail to translocate along Purkinje cell dendrites. Importantly, given that AP-2 is deleted only in Purkinje cells and not in the olivary neurons that give rise to the CFs, this loss of CF innervation is a direct result of PC dysfunction. Third, hyperexcitation is already observed in 8-day-old Purkinje cells cultured ex vivo for three weeks and can be rescued by upregulating synaptic glutamate reuptake using ceftriaxone. Since perturbation of postsynaptic activity of Purkinje cells has been previously shown to regulate synaptic translocation of CFs (Lorenzetto et al., 2009), we believe that hyperexcitation induced by the gain of PF synapses is a cause of decreased CF innervation in adult Purkinje cells. This hypothesis is also in line with published data indicating that CF elimination is prevented in spontaneously occurring mutant mouse strains, including *weaver* (Crepel and Mariani, 1976), *staggerer* (Crepel et al., 1980), and *reeler* (Mariani et al., 1977), in which the granule cells that give rise to PFs are not formed or degenerate. Consistently, our data indicate that deletion of AP-2 after postnatal day 21 (when the CF territory is already fully established) does not affect their translocation. CFs provide essential instructive signals for associative cerebellar learning (Silva et al., 2024). Changes in CF input occur in SCA (Barnes et al., 2011; Blake et al., 2013; Osório and White, 2023; Smeets and Verbeek, 2016) and genetic silencing of olivocerebellar synapses leads to severe motor deficit in mice (White and Sillitoe, 2017). Thus, we suggest that an almost complete loss of CF innervation as a consequence of increased Purkinje cell hyperexcitation underlies motor impairments in AP-2 cKO mice, a hypothesis that needs to be tested in future experiments.

Although our results are consistent with the idea that PF synaptogenesis increases Purkinje cell hyperexcitation leading to blockade of CF translocation in mice with conditional deletion of AP-2 in Purkinje cells, there are also alternative interpretations of our phenotype. An excess of PF inputs may provide a strong source of trophic support that could inhibit the translocation of CFs along Purkinje cell dendrites. Indeed, alterations in BDNF and its receptor tropomyosin kinase B (TrkB) have been recently reported in a mouse model of SCA6 (Cook et al., 2022; Cook et al., 2023), whereas inactivation of TrkB has been shown to slow CF elimination (Johnson et al., 2007). Likewise, synaptic transmission at PF-Purkinje cell synapses is largely mediated by AMPA-type glutamate receptors (Gutierrez-Castellanos et al., 2017). The overactivation of these receptors and downstream signal transduction cascades can therefore influence Purkinje cell metabolism in a way that has an inhibitory effect on CFs. Additionally, AP-2 may also contribute to synaptic plasticity by directly regulating AMPA receptor trafficking at dendritic spines. For instance, components of CME have previously been reported to regulate cerebellar LTD (Wang and Linden, 2000), whereas the AP-2α subunit has been shown to interact with the actin-based motor myosin-6 in Purkinje cells, and this interaction is crucial for LTD-induced endocytosis of AMPA receptors from the surface of dendritic spines (Wagner et al., 2019). Although, we did not detect changes in myosin-6 and/or AMPA receptors in the AP-2 cKO cerebellum in the current study, this interaction may still be required to recycle AMPA receptors during synaptic transmission and thereby also influence glutamate-induced excitotoxicity.

In summary, our study provides unprecedented insights into the molecular mechanisms underlying AP-2-mediated synaptic regulation in motor coordination. These findings hold significant implications for understanding the pathophysiology of ataxia and other cerebellar disorders linked to synaptic dysfunction. Moreover, our observations are relevant to a recently described mutation in *AP2M1* that has been associated with developmental encephalopathy accompanied by ataxia (Helbig et al., 2019). We propose that the ataxic symptoms observed in these patients may arise from disrupted cerebellar synaptogenesis attributable to AP-2 loss-of-function. Our restoration of cerebellar network excitability using ceftriaxone underscores the potential of targeting synaptic transmission as a promising therapeutic avenue for managing ataxia associated with cerebellar dysfunction.

## Data availability

All data needed to evaluate the conclusions in the paper are present in the paper and/or the source data. Proteome data of all experiments are deposited in the database PRIDE and/or Zenodo and publicly accessible after publishing. Source data are provided with this paper. Additional data related to this paper may be requested from the corresponding author.

## Acknowledgements

We thank S. Müller and Dr. M. Schröter for their expert assistance. We are indebted to Dr. C. Jüngst (CECAD Imaging Facility), Dr. S. Müller and Dr. J.-W. Lackmann (CECAD Proteomic Facility) for their help and expert assistance. We thank Prof. E. Rugarli (CECAD, University of Cologne) for providing L7-Cre transgenic mice and Prof. M. Bergami (CECAD, University of Cologne) for providing Ai9-tdTomato transgenic mice. The work of NLK is funded by the Deutsche Forschungsgemeinschaft (DFG, German Research Foundation): EXC 2030–390661388, KO 5091/4-1, DFG-431549029–SFB 1451, DFG-233886668-GRK1960 and DFG-411422114 - GRK 2550.

## Author contributions

Conceptualization: NLK

Methodology: MT, FL, AP, SV, GG, NLK

Investigation: MT, JT, EO, IK, EK, QS, SV, GG, NLK

Visualization: MT, IK, AP, SV, GG, NLK

Supervision: TK, GS, SV, GG, NLK

Writing—original draft: MT& NLK

Writing—review & editing: NLK

