## Supplementary material for "Endocytic adaptor AP-2 maintains Purkinje cell function by balancing cerebellar parallel and climbing fiber synapses"

**This PDF file includes:**

Supplementary figures 1-6

**STAR ★ Methods**

**Key Resources Table**

Legends for Videos S1-8

Legends for Table S1-5

**Supplementary figures 1-6.**

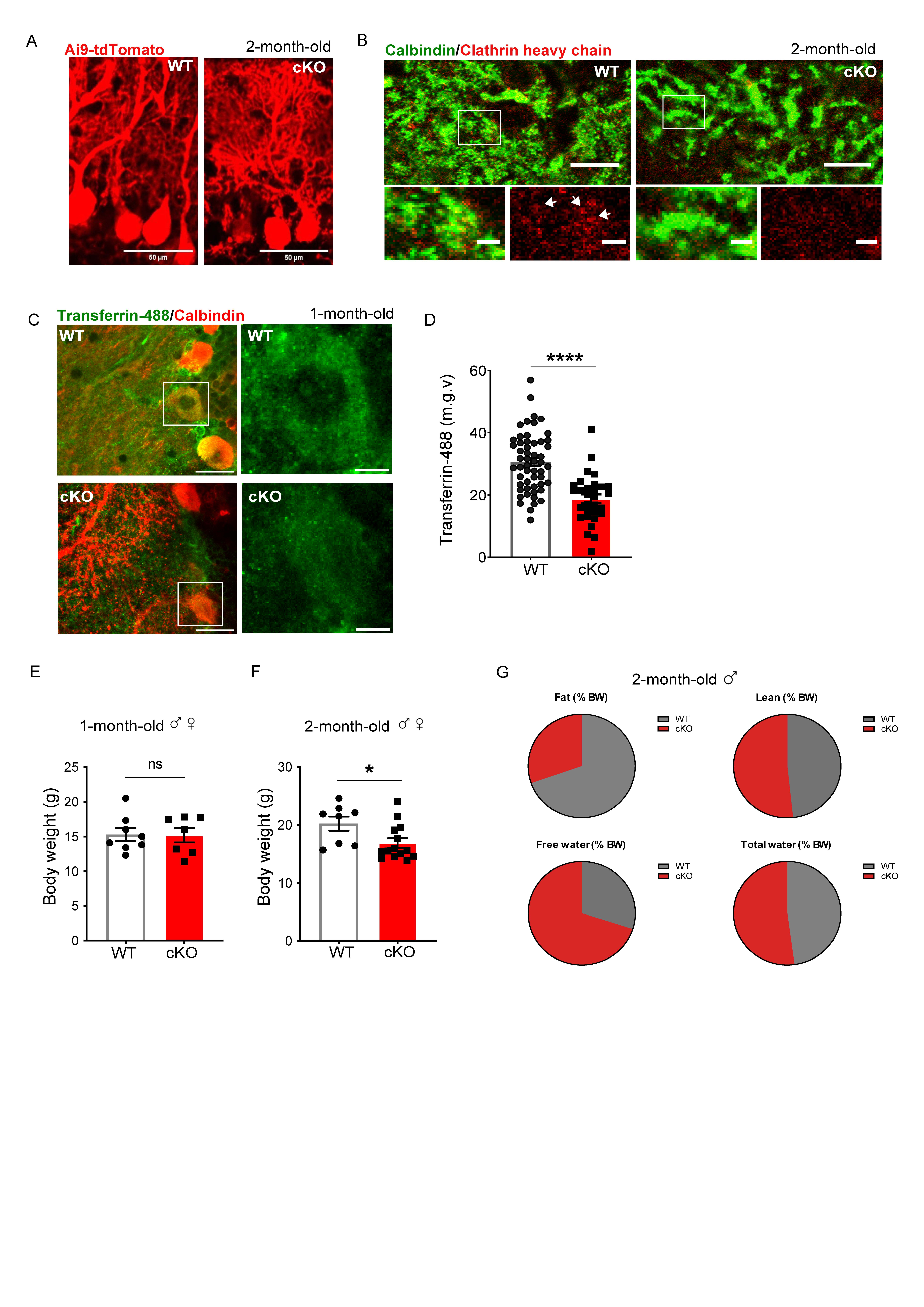

**Supplemental figure1. Related to Figure 1. Characterization of AP-2 cKO mice.**

**(A)** Immunofluorescence staining for tdTomato showing the *L7^Cre^* mediated recombination in cerebellum of 2-month-old WT and AP-2 cKO mice crossed with a reporter mouse line (Ai9-tdTomato). Scale bar, 50µm.

**(B)** Immunofluorescence staining for Calbindin (green) and Clathrin Heavy Chain (red) on sagittal cerebellar slices from 2-month-old WT and AP-2 cKO mice. Arrowheads indicate the presence of clathrin foci in WT Purkinje cells, whereas clathrin localization is dispersed in AP-2 cKO Purkinje cells. Scale bars: 20 µm, 2µm inserts.

(**C,D**) Analysis of Transferrin-488 uptake in acute cerebellar slices from 4-week-old WT and AP-2 cKO mice (WT: 30.57±1.286 m.g.v, cKO: 19.85±1.761 m.g.v., p<0.0001). Statistical significance was determined by unpaired two-tailed Student’s t-test.

**(E,F)** Body weight quantification in 1-month-old (C) and 2-month-old (D) WT and AP-2 cKO mice. For quantification, both sexes were pooled together. At two months of age, cKO mice display a significantly decreased body weight compared to WT. Each dot represents one mouse ((N=8 for WT, N= 7 for cKO, for 1 month of age, N=8 for WT, N= 14 for cKO for 2 months of age). Statistical significance was determined by unpaired two-tailed Student’s t-test (for 2 months of age, WT: 20g ± 1.2g, cKO: 17g ± 0.82g; p=0.0278).

**(G)** Body composition analysis in 2 months-old WT and AP-2 cKO male mice. Quantification of fat, lean, free water and total water as body weight percentage did not reveal any significant difference in body composition between WT (N=3) and cKO (N=4) mice. Statistical significance was determined by unpaired two-tailed Student’s t-test (Fat: WT: 8.914% ± 2.331%, cKO: 5.316% ± 0.5705%, p=0.1417; Lean: WT: 86.27% ± 2.101%, cKO: 87.66% ± 1.051%, p= 0.5482; Free water: WT: 0.9650% ± 0.8069%, cKO: 1.359% ± 0.4604%, p= 0.5050; Total water: WT: 77.17% ± 1.686%, cKO: 78.98% ± 1.098%, p= 0.3890).

Data is presented as mean±SEM. * p ≤0.05; ** p≤0.01; *** p≤0.001; **** p≤0.0001. n.s.-non-significant.

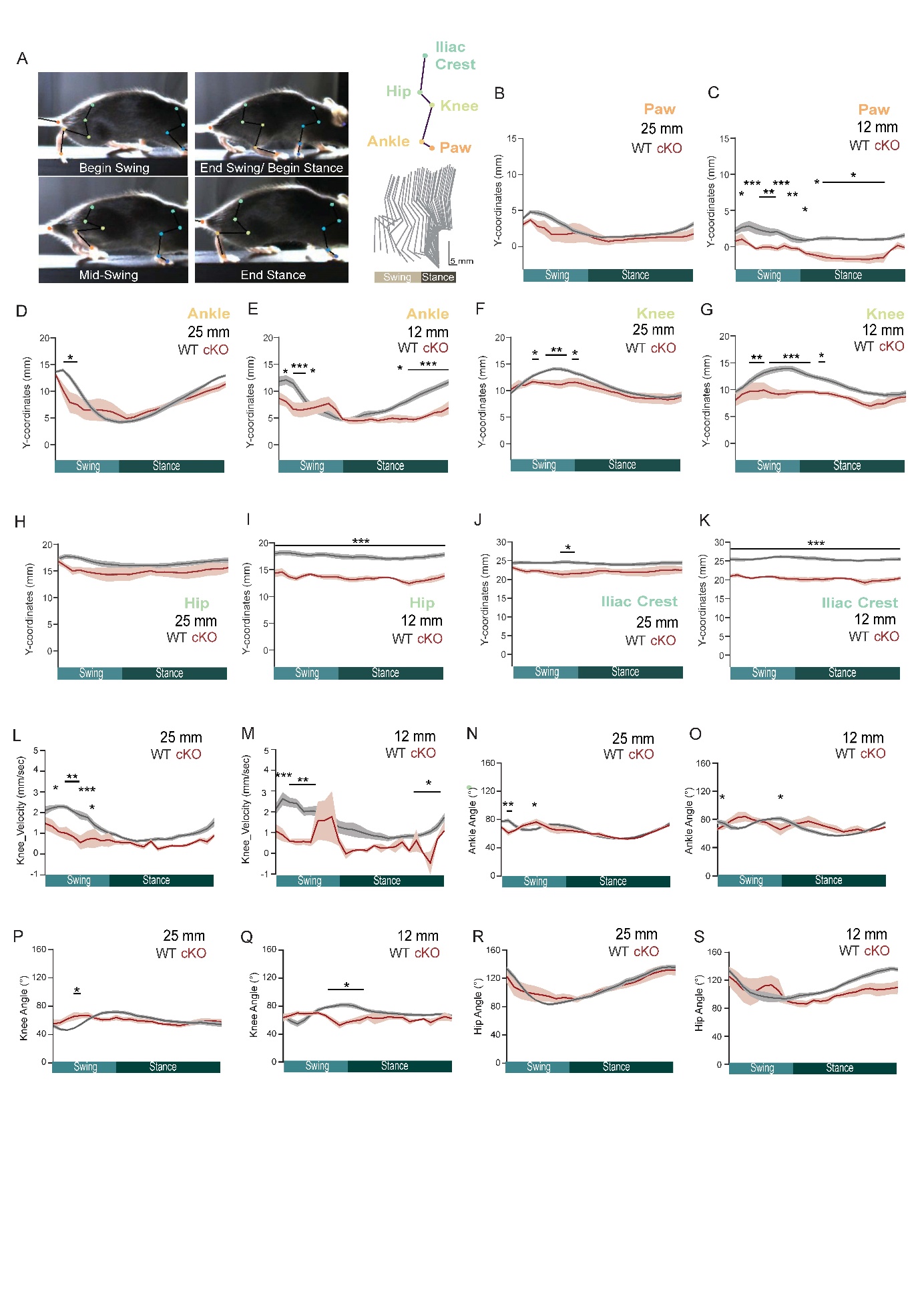

**Supplemental figure 2. Related to figure 1. Limb kinematic alterations in AP-2 cKO mice.**

**(A)** Image showing example of kinematic tracking during a step cycle.

(**B-K**) Line graphs showing a detailed analysis of y-coordinates (height) of the hindpaw (B,C), ankle (D,E), knee (F,G), hip (H,I) and iliac crest (J,K) in WT and AP-2 cKO mice walking on a 25mm-(B,D,F,H,J) or 12 mm-wide beam (C,E,G,I,K). Pooled data are shown in Fig. 1K-N. Statistical significance was determined by two-way ANOVA followed by Šidák multiple comparison between joints in WT and AP-2 cKO mice (see Table S4 for the detailed results of two-way ANOVA multiple comparison).

**(L,M)** Line graphs showing the variation in knee velocity during a normalized step cycle in WT and AP-2 cKO mice walking on a 25mm-(L) or 12 mm-wide beam (M). On the 25 mm beam, knee velocity was significantly altered during swing phase in AP-2 cKO mice compared to WT (L; N=9 for WT, N=6 for cKO). Additionally, the velocity was altered during stance on the 12 mm beam (M; N=9 for WT, N=6 for cKO). Statistical significance was determined by two-way ANOVA followed by Šidák multiple comparison between joints in WT and AP-2 cKO mice (see Table S4 for the detailed results of two-way ANOVA multiple comparison).

**(N-S)** Line graphs showing the variation in ankle, knee and hip angles during a normalized step cycle in WT and AP-2 cKO mice walking on a 25 mm (N,P,R; N=9 for WT, N=6 for cKO) or 12mm beam (O,Q,S; N=9 for WT, N=6 for cKO). The relative positions of the ankle and knee joint differ during swing phase in AP-2 cKO mice compared to WT while crossing a 25 mm (N,P) and 12 mm beam (O,Q). Statistical significance was determined by two-way ANOVA followed by Šidák multiple comparison between joints in WT and AP-2 cKO mice (see Table S4 for the detailed results of two-way ANOVA multiple comparison).

Data presented in Fig. S2 B-S show mean as a filled dark line, and SEM as the shaded area around it. * p ≤0.05; ** p ≤0.01; *** p ≤0.001.

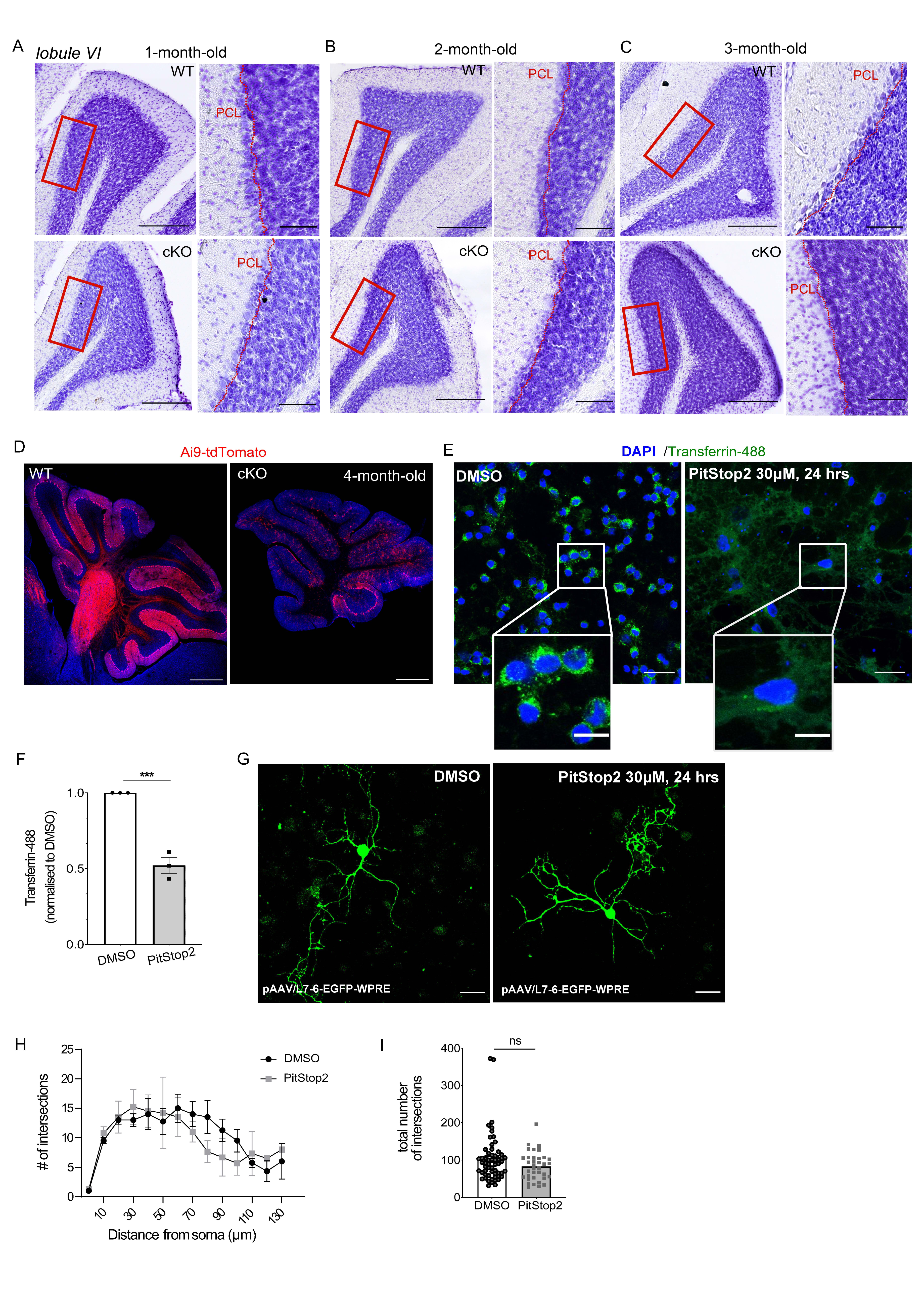

**Supplemental figure 3. Related to figure 2. The morphological changes of Purkinje cells in AP-2 cKO mice are independent of clathrin.**

**(A-C)** Nissl staining of cerebellar sagittal sections from WT and cKO mice at 1 month (A), 2 months (B) and 3 months (C) of age. Scale bars: 500 µm, inserts 100 µm.

(**D**) tdTomato expression in cerebellum of 4-month-old WT and AP-2 cKO mice crossed with a reporter mouse line (Ai9-tdTomato: *Rosa^loxP-STOP-loxP-tdtomato^*) shows an almost complete loss of Purkinje cell at this age. Scale bar, 500 µm.

**(E)** Transferrin-488 (green) uptake in DIV 19 cerebellar primary neurons treated with either DMSO (control) or the clathrin inhibitor PitStop2, and stained with DAPI (blue) to label nuclei. Scale bar: 25 µm, inserts 10µm.

**(F)** Quantification of Transferrin-488 uptake in WT DIV 19 cerebellar primary neuronal culture shows significant decrease in the uptake after PitStop2 treatment in comparison to DMSO set to 1. Each dot represents one cell (DMSO: n=565 cells from N=3 independent experiments, PitStop2: n=572 cells from N=3 independent experiments). Statistical significance was determined by one-tailed unpaired Student’s t-test (DMSO: 1 ± 0.015, PitStop2: 0.54 ± 0.0090; p<0.0001).

**(G)** Immunostaining for GFP on WT DIV 19 primary cerebellar PCs transduced with AAV2/rh10-L7-6-EGFP-WPRE and treated with either DMSO or PitStop2. Scale bar: 20 µm.

**(H,I)** Sholl analysis of the number of intersections from the cell soma to the most distal neurites of WT DIV 19 primary cerebellar PCs transduced with AAV2/rh10-L7-6-EGFP-WPRE and treated with either DMSO or PitStop2. No difference was observed in the number of intersections from most proximal to most distal distance from the soma (H, N=4 independent experiments), neither in the total number of intersections (I, n=58 cells and n==36 cells for DMSO and PitStop2 respectively from N=4 independent experiments). Statistical significance was determined by one-tailed unpaired Student’s t-test (DMSO: 102.4 ± 8.571, PitStop2: 82.67 ± 6.328; p=0.1027).

Data is presented as mean±SEM. * p ≤0.05; ** p≤0.01; *** p≤0.001; **** p≤0.0001. n.s.-non-significant.

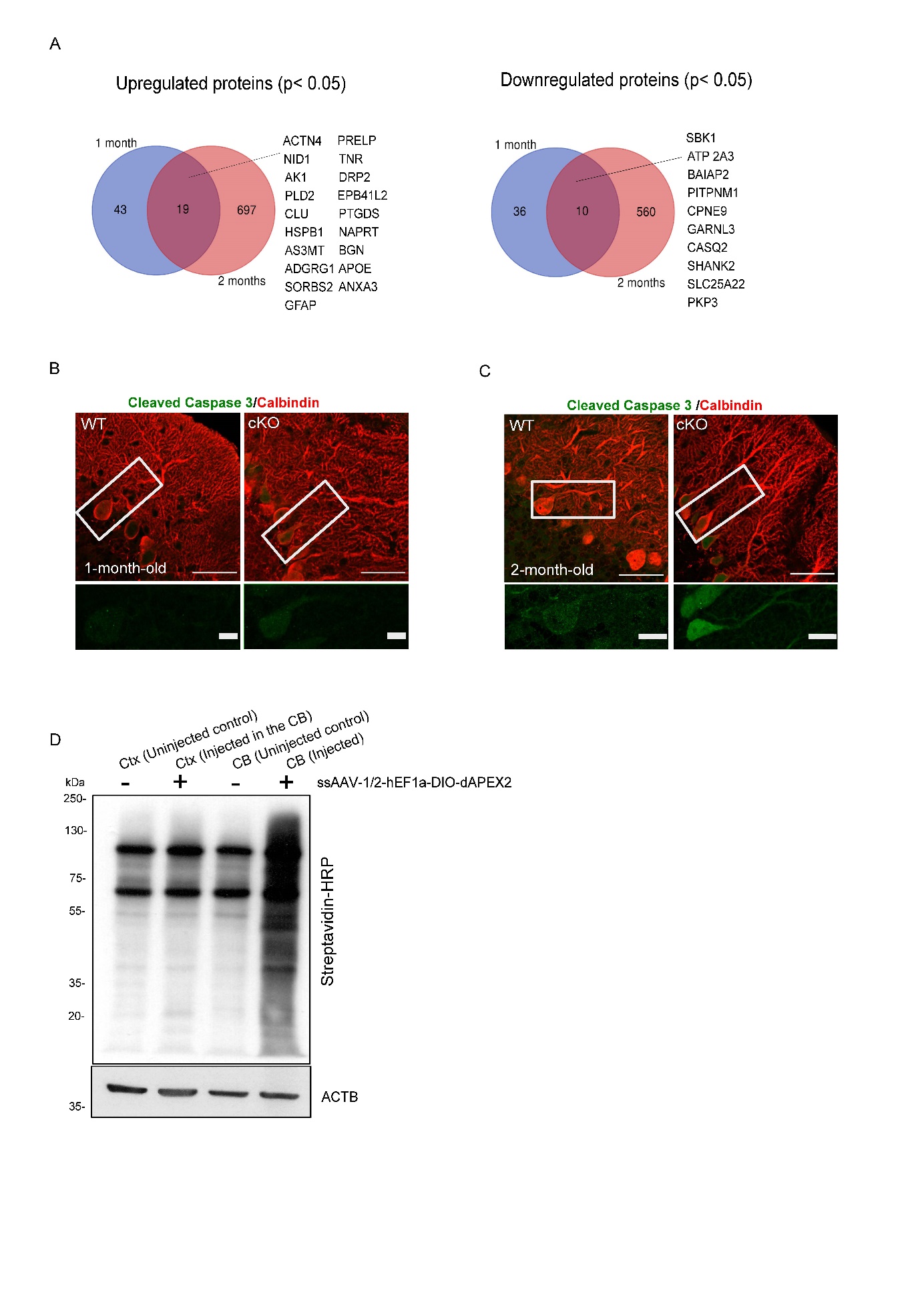

**Supplemental figure 4. Related to figure 3.**

(A) Venn diagrams showing commonly up- and downregulated proteins at p≤0.05 in the cerebellum of 1-month-old (left) and 2-month-old (right) AP-2 cKO mice (N=5 per genotype).

**(B,C)** Immunofluorescence staining for Calbindin (red) and Cleaved-caspase 3 (green) in the cerebellum of 1- month-old (B) and 2-month-old (C) WT and AP-2 cKO Purkinje cells. At 2 months of age, AP-2 cKO Purkinje cells displayed an increase in Cleaved-caspase 3 expression in line with proteomics data (see Fig. 3B). Scale bars: 50 µm, inserts 10 µm.

(**D**) Western blotting of cortical (Ctx) and cerebellar (CB) protein lysates from mice injected with ssAAV-1/2-hEF1a-DIO-dAPEX2 and controls stained with streptavidin-horseradish peroxidase after streptavidin bead enrichment in the presence of biotin phenol and H2O2. Each lane includes endogenously biotinylated proteins at around 70 and 100 kDa.

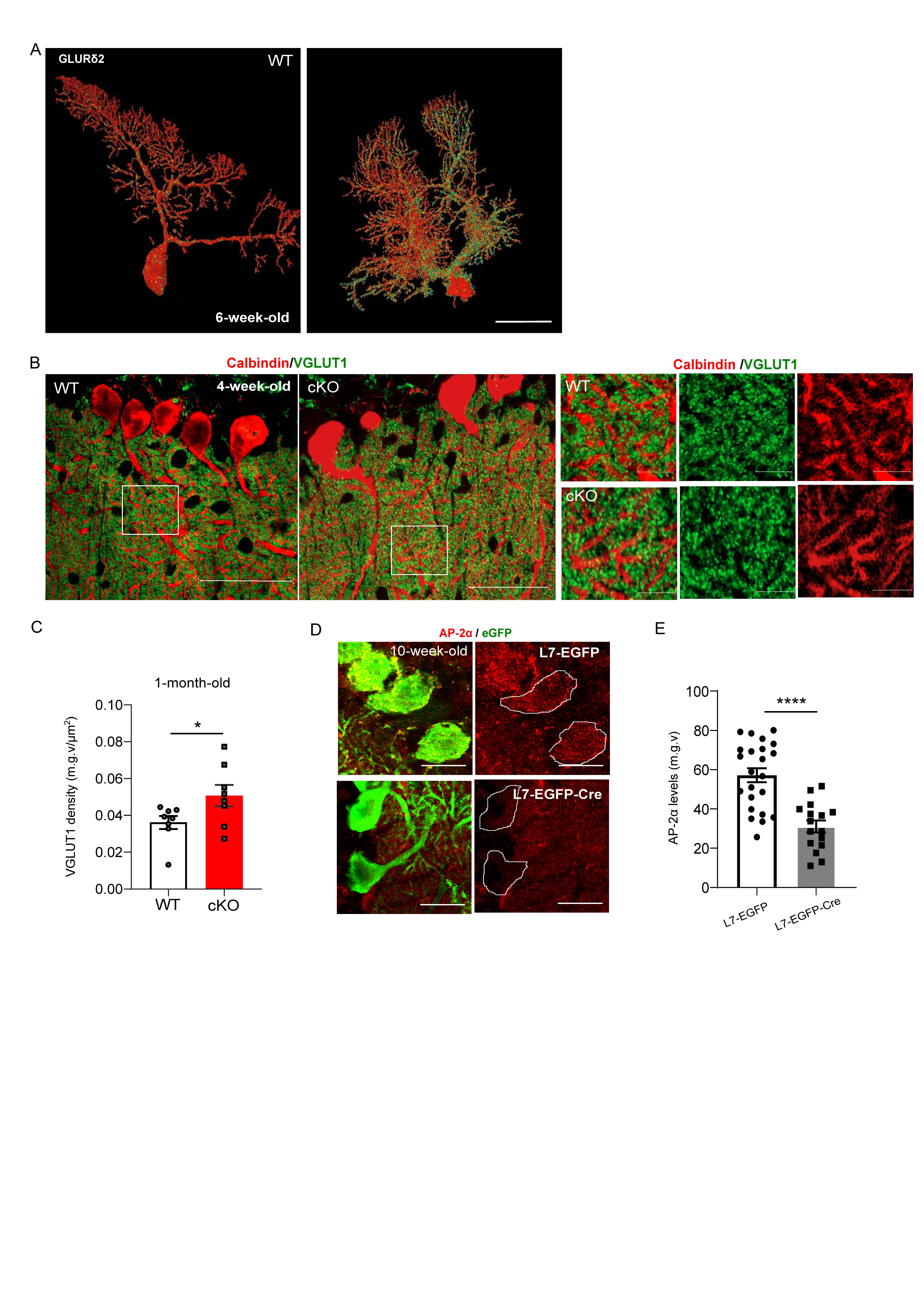

**Supplemental figure 5. Related to figure 5 and 6.**

**(A)** AMIRA-based 3D visualization of GLURδ2 distribution in 6-week-old Purkinje cells of WT and cKO mice. 3D reconstruction of Purkinje cells is based on GFP immunostaining after transduction with AAV1/2-Ef1α-DIO-EYFP. Scale bar: 50µm.

**(B)** Representative confocal images of 1-month-old WT and AP-2 cKO cerebellum stained with antibodies against Calbindin (red) to label Purkinje cells and VGLUT1 (green) to label PF synapses. Scale bar: 50µm, inserts 20µm.

**(C)** Quantification of VGLUT1 density in 4-week-old WT and AP-2 cKO Purkinje cells. Each dot represents one image (WT/cKO 8 images from N=3 mice). Statistical significance determined by unpaired two-tailed Student’s t-test (WT: 0.036 ± 0.003, cKO: 0.051 ± 0.006; p=0.0493).

(**D**) Representative confocal images of 10-week-old Purkinje cells transduced with either AAV2/rh10-L7-6-EGFP-P2A-Cre-WPRE (resulting in acute AP-2 cKO) or control virus AAV2/rh10-L7-6-EGFP-WPRE. Scale bar: 20µm.

**(E)** Analysis of AP-2α levels in Purkinje cells from *Ap2m1^flox/flox^* mice injected with either AAV AAV2/rh10-L7-6-EGFP-WPRE control virus (WT) or AAV2/rh10-L7-6-EGFP-P2A-Cre-WPRE (cKO). Each dot represents one cell (WT n=23 and cKO n=16 cells from N=3 mice). Statistical significance determined by unpaired two-tailed Student’s t-test (WT: 57.12 ± 3.547, cKO: 31.08 ± 3.038; p<0.0001).

Data is presented as mean±SEM. * p ≤0.05; ** p≤0.01; *** p≤0.001; **** p≤0.0001.

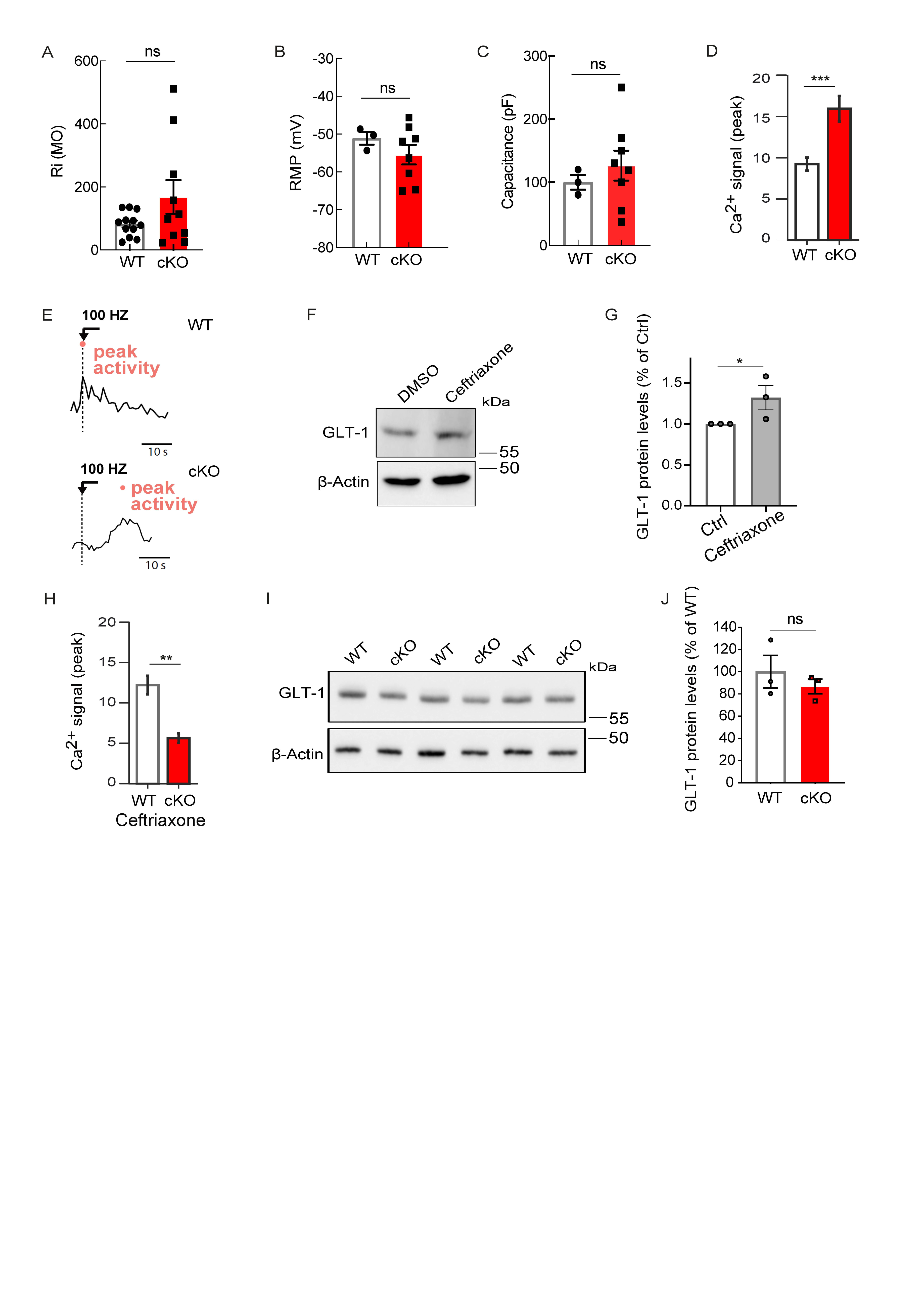

**Supplemental figure 6. Related to figure 7. No alterations in intrinsic properties of AP-2 cKO Purkinje cells compared to WT.**

**(A-C)** Analysis of Purkinje cell intrinsic properties in 4 weeks old WT and AP-2 cKO mice. No significant changes were detected in cKO Purkinje cell membrane resistance (A; WT n=12 cells from N=6 mice, cKO n=10 cells from N=3 mice), resting membrane potential (B; WT n=3 cells from N=3 mice, cKO n=8 cells from N=2 mice) and capacitance (C; WT n=3 cells from N=2 mice, cKO n=8 cells from N=2 mice). Statistical significance was determined by two-tailed Mann-Whitney test. The raw data of the patch clamp recordings are listed in Table S5. (**D**) Amplitude of somatic Ca^2+^ transients in response to 100 Hz stimulation in Purkinje cells transfected with ssAAV-9/2-mCaMKIIα-jGCaMP7f-WPRE-bGHp(A) in WT and AP-2 cKO cerebellar organotypic slices (WT: 9.233±0.432, cKO: 15.934±0.10, p<0.0001). Data are from WT (grey, n = 457 cells) or cKO (red, n = 255 cells) cerebellar organotypic slices (from N=6 WT and N=5 cKO mice). Statistical significance was determined by Welch´s t-test.

(**E)** Exemplary time point of maximal stimulus-evoked Ca2+ responses in WT and AP-2 cKO Pukinje cells upon their stimulation at 100Hz. Red dot indicates time point of peak response used for analysis in Figs 7J.

(**F,G**) Immunoblot analysis of GLT-1 levels in lysate obtained from WT DIV21 cerebellar organotypic cultures treated for 7 consecutive days with ceftriaxone. β-actin was used as loading control (WT set to 1, KO 1.321±0.151, p=0.049). Statistical significance was determined by one-tailed unpaired Student’s t-test.

(**H**) Amplitude of stimulus-evoked responses after 100µM ceftriaxone treatment (7 days starting at DIV15) in WT and AP-2 cKO Purkinje cells transfected with ssAAV-9/2-mCaMKIIα-jGCaMP7f-WPRE-bGHp(A) (WT: 12.233±0.735, cKO: 5.661±0.442, p=0.001). Data are from WT (grey, n = 277 cells) or cKO (red, n = 164 cells) cerebellar organotypic slices (from N=4 WT and N=2 cKO mice). Statistical significance was determined by Welch´s t-test.

**(I,J)** Immunoblot analysis of GLT-1 levels in cerebellum of 2-month-old WT and AP-2 cKO mice. β-actin was used both as a loading control and for normalisation of protein levels. Each dot in (I) represents one mouse (N=3 per genotype). Statistical significance was determined by one-tailed unpaired Student’s t-test.

Data is presented as mean±SEM. * p ≤0.05; ** p≤0.01; *** p≤0.001; **** p≤0.0001. n.s.-non-significant.

**STAR ★ Methods**

**Key Resources Table**

| **REAGENT or RESOURCE** | **SOURCE** | **IDENTIFIER** |
| --- | --- | --- |
| **Antibodies** | | |
| mouse anti-AP-2α | BD Biosciences | Cat# 610501; RRID: [AB_397867](https://www.antibodyregistry.org/AB_397867) |
| mouse anti-AP-2µ | BD Biosciences | Cat# 611350, RRID: [AB_398872](https://www.antibodyregistry.org/AB_398872) |
| mouse anti-β actin | Sigma-Aldrich | Cat# A-5441, RRID: [AB_476744](https://www.antibodyregistry.org/AB_476744) |
| chicken anti-Calbindin | Novus Biologicals | Cat# NBP2-50028, RRID: [AB_2938765](https://www.antibodyregistry.org/AB_2938765) |
| mouse anti-clathrin heavy chain | homemade | N/A |
| rabbit anti-Cleaved Caspase3 | Cell Signaling | Cat# 9661S, RRID: [AB_2341188](https://www.antibodyregistry.org/AB_2341188) |
| chicken anti-GFP | Abcam | Cat#ab13970, RRID: [AB_300798](https://www.antibodyregistry.org/AB_300798) |
| rabbit anti-GLT-1 | Santa Cruz Biotechnology | Cat#sc-365634, RRID: [AB_10844832](https://www.antibodyregistry.org/AB_10844832) |
| rabbit anti-GLURδ2 (for WB) | Abcam | Cat# ab190358, RRID: N/A |
| rabbit anti-GLURδ2 (for IHC) | Novus Biologicals | Cat# NBP2-31723, RRID: N/A |
| rabbit anti-GRID1IP | Novus Biologicals | Cat# NBP1-94174, RRID: [AB_11008710](https://www.antibodyregistry.org/AB_11008710) |
| guinea pig anti-Vglut1 | Synaptic Systems | Cat# 135304, RRID: [AB_887878](https://www.antibodyregistry.org/AB_887878) |
| guinea pig anti-Vglut2 | Synaptic Systems | Cat# 135404, RRID: [AB_887884](https://www.antibodyregistry.org/AB_887884) |
| goat anti-chicken Alexa 488 | Thermo Fisher Scientific | Cat# A11039, RRID: [AB_2534096](https://www.antibodyregistry.org/AB_2534096) |
| goat anti-chicken Alexa 568 | Thermo Fisher Scientific | Cat# A11041, RRID: [AB_2534098](https://www.antibodyregistry.org/AB_2534098) |
| goat anti-chicken Alexa 647 | Thermo Fisher Scientific | Cat# A21449, RRID: [AB_2535866](https://www.antibodyregistry.org/AB_2535866) |
| goat anti-guinea pig Alexa 488 | Thermo Fisher Scientific | Cat# A11073, RRID: [AB_2534117](https://www.antibodyregistry.org/AB_2534117) |
| goat anti-guinea pig Alexa 647 | Thermo Fisher Scientific | Cat# A21450, RRID: [AB_2535867](https://www.antibodyregistry.org/AB_2535867) |
| goat anti-mouse Alexa 488 | Thermo Fisher Scientific | Cat# A11029, RRID: [AB_2534088](https://www.antibodyregistry.org/AB_2534088) |
| goat anti-mouse Alexa 568 | Thermo Fisher Scientific | Cat# A11031, RRID: [AB_144696](https://www.antibodyregistry.org/AB_144696) |
| goat anti-mouse Alexa 647 | Thermo Fisher Scientific | Cat# A21236, RRID: [AB_2535805](https://www.antibodyregistry.org/AB_2535805) |
| goat anti-rabbit Alexa 488 | Thermo Fisher Scientific | Cat# A11034, RRID: [AB_2576217](https://www.antibodyregistry.org/AB_2576217) |
| goat anti-rabbit Alexa 568 | Thermo Fisher Scientific | Cat# A11011, RRID: [AB_143157](https://www.antibodyregistry.org/AB_143157) |
| goat anti-rabbit Alexa 647 | Thermo Fisher Scientific | Cat# A21245, RRID: [AB_2535813](https://www.antibodyregistry.org/AB_2535813) |
| Goat anti-Rabbit IgG (H+L) peroxidase-conjugated | Sigma-Aldrich | Cat# A0545, RRID: [AB_257896](https://www.antibodyregistry.org/AB_257896) |
| Rabbit anti-Chicken IgG (H+L) peroxidase-conjugated | Merk Millipore | Cat# AP162P, RRID: [AB_91653](https://www.antibodyregistry.org/AB_91653) |
| Rabbit anti-Mouse IgG (H+L) peroxidase-conjugated | Sigma-Aldrich | Cat# A9044, RRID: [AB_258431](https://www.antibodyregistry.org/AB_258431) |
| Normal rabbit IgG | Cell Signaling | Cat#2729S, RRID: AB_1031062 |
| **Viruses** | | |
| AAV1/2-Ef1α-DIO EYFP | Karl Deisseroth (unpublished) | Addgene viral prep #27056-AAV1, RRID: [Addgene_27056](https://www.addgene.org/27056/) |
| AAV2/rh10-L7-6-EGFP-WPRE | custom-produced, this paper | To be deposited on Addgene upon publication: https://www.addgene.org/Günter_Schwarz/ |
| AAV2/rh10-L7-6-EGFP-P2A-Cre-WPRE | custom-produced, this paper | To be deposited on Addgene upon publication: https://www.addgene.org/Günter_Schwarz/ |
| ssAAV-9/2-mCaMKIIα-jGCaMP7f-WPRE-bGHp(A) |  |  |
| **Chemicals, peptides, and recombinant proteins** | | |
| Ascorbic acid | Carl Roth | 3525.1 |
| ß-mercaptoethanol | Carl Roth | 4227.1 |
| bis-tris-propane | Sigma Aldrich | B6755 |
| Bovine serum albumin (BSA) | Sigma Aldrich | A7906 |
| Bromophenol (0.03%) | Sigma Aldrich | B5525 |
| BSA (fatty acid free) | Sigma Aldrich | A6003 |
| Calcium chloride (CaCl_2_) | Carl Roth | 5239.2 |
| Ceftriaxone disodium salt hemi(heptahydrate) | Sigma Aldrich | C5793 |
| D-Glucose | Sigma Aldrich | G5767 |
| Dimethyl sulfoxide (DMSO) | Carl Roth | A994.2 |
| DMEM | Thermo Fisher Scientific | A14430 |
| dPBS | Gibco | 14190250 |
| DNAse | Sigma Aldrich | 150000U |
| EBSS | Thermo Fisher Scientific | 14155-048 |
| Fetal bovine serum (FBS) | Merck Millipore | S0115 |
| Gelatin from porcine skin | Sigma Aldrich | G2500 |
| GlutaMAX^TM^ | Thermo Fisher Scientific | 35050-061 |
| Glycerol | Carl Roth | 7530.1 |
| HBSS | Gibco | 14175-053 |
| Heparin | Sigma Aldrich | H4784 |
| HEPES | Thermo Fisher Scientific | 15630-080 |
| Horse serum (heat-inactivated) | Gibco | 26050088 |
| IGEPAL CA-630 | Sigma Aldrich | I8896 |
| Ketamin hydrochloride | Sigma Aldrich | K2753 |
| Kynurenic acid | Tocris | 3694 |
| Magnesium-ATP (Mg-ATP) | Sigma Aldrich | 20-113 |
| Magnesium chloride (MgCl_2_) | Carl Roth | 2189.1 |
| Magnesium sulfate (MgSO_4_) | Carl Roth | T888.1 |
| MEM | Sigma Aldrich | 7278 |
| Penicillin/Streptomycin (P/S) | Thermo Fisher Scientific | 15140-122 |
| Phosphocreatine | Tocris | 4325 |
| Pitstop | Abcam | ab120687 |
| Potassium chloride (KCl) | Karl Roth | 6781.1 |
| Potassium-gluconate | Merk Millipore | 299-27-4 |
| Protease and Phosphatase inhibitor mini tablets | Thermo Fisher Scientific | A32959 |
| Rompun 2% (Xylazin) | Bayer | KP0BZPE |
| Saponin | Sigma aldrich | 47036 |
| SDS | Carl Roth | 2326.2 |
| Sodium bicarbonate (NaHCO_3_) | Carl Roth | 8551.1 |
| Sodium chloride (NaCl) | Carl Roth | 3957.1 |
| Sodium deoxycholate | Thermo Fisher Scientific | 89904 |
| Sodium-GTP (Na-GTP) | Merk Millipore | G3776 |
| Sodium hydrogen phosphate (NaH_2_HPO_4_) | Carl Roth | 3904.1 |
| Sodium pyruvate | Thermo Fisher Scientific | 11360-039 |
| Soybean Trypsin Inhibitor | Merck Millipore | 10109886001 |
| Transferrin from human serum, Alexa Fluor 488 conjugate | Thermo Fisher Scientific | T13342 |
| Tris | VWR | 28.808.294 |
| Tris-HCl | Sigma Aldrich | T3253 |
| Triton X-100 | Sigma Aldrich | 9036-19-5 |
| Tween 20 | VWR | 663684B |
| Trypsin | Sigma Aldrich | T1005 |
| **Critical commercial assays** | | |
| RNAScope® Fluorescent Multiplex Detection Reagents | Advanced Cell Diagnostic | 323110 |
| *Mm-Grid2ip*-C1 RNA probe | Advanced Cell Diagnostic | Cat# 1253061-C1 |
| NEBuilder HiFi DNA Assembly | New England Biolabs | Cat# E2621S |
| **Deposited data** | | |
| Imaging and quantitative data | Mendeley data |  |
| Proteomics data | Mendeley data |  |
| Calcium imaging data | Mendeley data |  |
| **Experimental models: Organisms/strains** | | |
| *Ap2m1^flox^* | Prof. Dr. Volker Haucke, The Leibniz-Forschungsinstitut für Molekulare Pharmakologie, Berlin | Kononenko et al., 2014 |
| *L7^Cre^* (provided by Prof. Rugarli, Cologne) | The Jackson Laboratory | RRID: MGI: J:66884 |
| *B6.Cg-Gt(ROSA)26Sortm14(CAG-tdTomato)Hze (Ai9-tdTomato* (provided by Prof. Bergami, Cologne) | The Jackson Laboratory | RRID: MGI: J:155793 |
| **Oligonucleotides** | | |
| Primers for Ai9, see Table S3 | The Jackson Laboratory | N/A |
| Primers for *Ap2m1^flox^*, see Table S3 | Kononenko et al., 2014 | N/A |
| Primers for L7^Cre^, see Table S3 | The Jackson Laboratory | N/A |
| **Recombinant DNA** | | |
| Plasmid: pAAV/L7-6-GFP-WPRE | Nitta et al., 2017 | Addgene plasmid # 126462, RRID: [Addgene_126462](https://www.addgene.org/126462/) |
| Plasmid: pAAV-hSyn-mScarlet | Marshel et al.,2019 | Addgene plasmid #131001, RRID:[Addgene_131001](https://www.addgene.org/131001/) |
| Plasmid: pAAV.CMV.HI.eGFP-Cre.WPRE.SV40 | James M. Wilson (unpublished) | Addgene plasmid # 105545, [RRID:Addgene_105545](https://www.addgene.org/105545/) |
| Plasmid: pAdDeltaF6 | James M. Wilson (unpublished) | Addgene plasmid #112867, [RRID:Addgene_112867](https://www.addgene.org/112867/) |
| pAAV2/rh10 | James M. Wilson (unpublished) | Addgene plasmid # 112866, [RRID:Addgene_112866](https://www.addgene.org/112866/) |
| **Software and algorithms** | | |
| Amira Software 2020.2 Thermo Fisher Scientific | Thermo Fisher Scientific | <http://www.fei.com/software/amira-3d-for-life-sciences/>  RRID: SCR_007353 |
| Aperio ImageScope version 12.4.3.5008 | Leica Microsystems | <https://www.leicabiosystems.com/aperio-imagescope/>  RRID: SCR_020993 |
| pClamp | Molecular Devices | <http://www.moleculardevices.com/products/software/pclamp.html>  RRID: SCR_11323 |
| Excel – Office 2021 | Microsoft | <https://www.microsoft.com/>  RRID: SCR_016137 |
| Fiji version 1.53 | Wayne Rasband, National Institute of Health, Bethesda, USA | [https://imagej.net](https://imagej.netD)/ RRID: SCR_002285 |
| GraphPad Prism version 9.5.1 | GraphPad | <http://www.graphpad.com/>  RRID: SCR_002798 |
| Inkscape 1.3.2 | Inkscape | <https://inkscape.org/>  RRID: SCR_014479 |
| LAS X Life Science Microscope Software | Leica Microsystems | <https://www.leica-microsystems.com/products/microscope-software/p/leica-las-x-ls/>  RRID: SCR_013673 |
| R | The R Foundation of Statistical Computing, Vienna, AT | [https://www.r-project.org/foundation](https://www.r-project.org/foundation/)  RRID: SCR_000432 |
| AutoGaitA | Hosseini et al. BioRxiv 2024 | https://github.com/mahan-hosseini/AutoGaitA |

**Resource availability**

*Lead contact*

**Materials availability**

pAAV/L7-6-EGFP-WPRE or pAAV/L7-6-EGFP-P2A-Cre-WPRE (<https://www.addgene.org/Gunter_Schwarz>).

**Experimental model and subject details**

Mice were kept on a C57BL6 background. Postnatal stages are indicated in Results and Figure legends. Mice were group-housed in polycarbonate cages with 12hr day/light cycles and water and food were available *ad libitum*. All experiments were performed in accordance with the regulations issued by the Federal Government of Germany, European Union legislation and the regulations of the University of Cologne. The experimental procedures were approved by the Landesamt für Natur, Umwelt und Verbraucherschutz Nordrhein-Westfalen (Permit Number: AZ 81-02.04.2020.A418, AZ 81-02.04.2021.A067, AZ 81-02.04.2021.A132 and AZ 81-02.04.2022.A116). *Ap2m1^flox^* mice have been previously described (Kononenko et al., 2014) and were crossed in this study with *L7^Cre^* mice (Barski et al., 2000) to generate AP-2 cKO mice (Genotype: *L7^Cre/+^;Ap2m1^flox/flox^*). In a subset of AP-2 cKO mice, the tdTomato expression was driven by crossing *L7^Cre/+^;Ap2m1^flox/flox^* ^mice^ with the reporter mouse line Ai9 (Madisen et al., 2010). For comparison of AP-2 cKO and control mice, littermates from several litters were used. Both female and male mice were used for primary neurons culture, cerebellar acute slices, cerebellar organotypic slices, immunostaining, immunoblotting, multiplex RNA in situ analysis, behavioural experiments, proteomics, electrophysiology and stereotactic viral vector injections. Only male mice were used for MRI body composition measurements. C57BL6/NRj mice were used for MS and Co-IPs WT experiments.

**Method details**

**Tissue processing and immunohistochemical analysis**

Adult mice were anaesthetised with an intraperitoneal injection of Ketanest/Rompun and transcardially perfused with Ringer solution (0.85% NaCl, 0.025% KCl, 0.02% NaHCO_3_, 0.01% heparin, pH 6.9) followed by 4% PFA in PBS (pH 7.4). Brains were dissected and post-fixed in 4% PFA overnight at 4°C and subsequently placed in a mixture of 20% (vol/vol) glycerol and 2% (vol/vol) dimethyl sulfoxide (VWR international) in 0.4M PBS for cryoprotection. 40 µm sagittal cryosections were obtained and free-floating sections were collected in the same cryoprotective solution mentioned above and stored at -80°C until further use.

For immunofluorescent staining, brain sections were washed once in PBS for 10 min and incubated in blocking solution containing 10% normal goat serum (NGS) in PBS plus 0.5% Triton X-100 (0.5% PBT) for 1 hr at room temperature (RT). Sections were incubated with primary antibodies for 48hr at 4°C in 0.3% PBT plus 3% NGS and afterwards washed 3 times for 5-10 min each in 0.3% PBT. Sections were then incubated with secondary antibodies for 2hr in the dark at RT in 0.3% PBT plus 3% NGS and washed 3 times for 5-10 min each in 0.3% PBT. Sections were finally mounted on gelatin-coated glass slides with Immu-Mount (Epredia). A list of primary and secondary antibodies is provided in the Key Resource Table. The following primary antibodies dilutions were used: 1:1000 for chicken anti-Calbindin, 1:1000 for chicken anti-GFP, 1:300 for rabbit anti- Cleaved Caspase3, 1:200 for mouse anti-AP-2α, 1:200 for rabbit anti-GRID2IP, 1:200 for rabbit anti-GLURδ2, 1:1000 for guinea pig anti-Vglut2 and 1:500 for guinea pig anti-Vglut1. All Alexa fluorophore-conjugated secondary antibodies were used at a 1:500 dilution.

**Nissl staining**

For Cresyl-violet staining 40 µm sagittal sections from 1-, 2- and 3-month-old AP-2 cKO mice and control littermates were mounted on SuperFrost Ultra plus microscope slides (Thermo Scientific) and dried overnight. Sections were stained following previously described protocol (Kononenko et al., 2017).

**Immunoblotting analysis**

2-month-old mice were sacrificed via cervical dislocation. Brains were isolated and the cerebellum was dissected, shock-frozen in liquid nitrogen and stored at -80°C until tissue lysis. For immunoblot analysis of cerebellar organotypic cultures, slices were collected at DIV21, shock-frozen in liquid nitrogen and stored at -80°C until tissue lysis. Samples were homogenized in RIPA buffer (50 mM Tris pH 8.0, 150 mM NaCl, 1.0% IGEPAL CA‐630, 0.5% Sodium deoxycholate, 0.1% SDS) containing phosphatase inhibitor (Thermo Scientific) and protease inhibitor (Roche) using a Wheaton Potter-Evehjem Tissue Grinder. Subsequently, samples were sonicated, incubated on ice for 45 min and centrifuged at 13000 rpm for 15 min at 4°C. Supernatant concentration was assessed using Bradford assay (Sigma) and samples were mixed with 4x SDS buffer (250 mM Tris-HCL, 1% (w/v) SDS, 40% (v/v) Glycerol, 4% (v/v) ß-mercaptoethanol, 0.03% Bromophenol) and boiled for 5 min at 95°C. 10-20 μg of protein per sample were loaded onto SDS-page gels for protein separation and afterwards transferred onto nitrocellulose membrane via full-wet transfer assay (Biorad). Membranes were blocked in 5% milk or bovine serum albumin (BSA) in TBS (20 mM Tris pH = 7.6, 150 mM NaCl) containing 1% Tween (TBS-T) at RT for 1h before incubation with primary antibodies in TBS overnight at 4°C. Afterwards, membranes were washed 3 times with TBS-T for 10 min at RT. before incubation with HRP-tagged secondary antibodies for 1h at RT. Membranes were finally washed 3 times with TBS-T for 10 min at RT. A list of primary and secondary antibodies is provided in the Key Resource Table. Protein levels were visualized using ECL-based autoradiography film system (Super RX-N, Fujifilm) or ChemiDocTM Imaging system (BioRad) and analyzed using Gel Analyzer plugin from ImageJ (Fiji). Protein levels were first normalized to loading control and to the appropriate control. The following primary antibodies dilutions were used: 1:1000 for mouse anti- AP-2α, 1:1000 for mouse anti-AP-2µ, 1:1000 for chicken anti-Calbindin, 1:1000 for rabbit anti-GRID2IP, 1:1000 for rabbit anti-GLT-1 and 1:2500 for mouse anti- β actin.

**Co-immunoprecipitation**

For immunoprecipitation experiments, 20 µL Dynabeads Protein G (Thermo Fischer Scientific) were coated with 2µg antibody targeting the protein of interest and corresponding IgG as a negative control (see Key Resource Table). Dynabeads storing solution was replaced with 100 µl PBS and 2µg of antibody was added. The beads were incubated with the antibody for 2-3h at 4°C on a shaker and then washed with 200µl PBS to remove excessive antibody. 8-week-old WT mice were sacrificed via cervical dislocation, brains were isolated and the cerebellum was dissected and homogenized in Co-IP buffer (50 mM Tris‐HCl pH = 7.4, 1% NP‐40/Igepal, 100 mM NaCl, 2 mM MgCl_2_) supplemented with Proteinase Inhibitor (Roche) und Phosphatase Inhibitor (ThermoScientific) using a Wheaton Potter‐Elvehjem Tissue Grinder. Samples were sonicated and incubated on ice for 45 min before being centrifuged at 13000 rpm for 20 min at 4°C. Protein concentration was assessed using Bradford assay (Sigma). An equal amount of protein was added to the antibody-coupled Dynabeads and control IgG for overnight incubation at 4°C on a shaker. Afterwards, the lysates were removed and Dynabeads were washed 3 times with Co-IP buffer before being dissolved in a mixture of 20 µL Co-IP buffer and 20 µL 4x SDS buffer and boiled at 95°C for 5 min. Precipitation of proteins was detected via SDS-page gel.

**Mass spectrometry (MS) analysis of AP-2α binding partners in the cerebellum**

8 weeks old WT mice were sacrificed via cervical dislocation. Brains were isolated and the cerebellum was dissected for MS analysis. Cerebellar tissue was homogenized in Co-IP buffer, as described in the previous section. Samples were boiled at 95 °C for 5 min and then loaded onto SDS-PAGE gels, reduced (DTT), and alkylated (CAA). Digestion was performed using trypsin at 37 °C overnight. Peptides were extracted and purified using Stagetips. Eluted peptides were dried in vacuo, resuspended in 1% formic acid/4% acetonitrile and stored at −20 °C before MS measurement. All samples were analyzed by the CECAD proteomics facility as previously described (Camblor-Perujo, Özer-Yildiz et al., 2023).

**Proteomics**

For total proteome analysis 2 and 3 months old mice were sacrificed via cervical dislocation. Brains were extracted and the cerebellum was dissected, shock frozen in liquid nitrogen and stored at -80°C until further use. Cerebellar tissue was lysed in Urea lysis buffer (50 mM TEAB, 8M Urea, 50x Protease inhibitor), sonicated and centrifuged at 20000g for 15 min. Protein concentration was assessed using Bradford assay (Sigma). Samples were processed with in-solution digestion. All solutions were provided by the CECAD proteomics facility. For each sample, 50 µg of protein were transferred into fresh tubes, reduced with 5 mM DTT for 1h at 25°C and subsequently alkylated with 40 mM CAA for 30 min in the dark. Protein digestion was performed by incubating samples in LysC at an enzyme:substrate ratio of 1:75 for 4h at 25°C. Samples were afterwards diluted with 50 mM TEAB to achieve a final concentration of 2 M Urea and then incubated overnight at 25°C in 1:75 ratio Trypsin. The following day, samples were acidified with formic acid (final concentration 1%). Peptides were extracted and purified using Stagetips. First, StageTips were equilibrated with washes in methanol, buffer B (80% acetonitril; 0.1% (v/v) formic acid) and twice buffer A (dH_2_O; 0.1% (v/v) formic acid). Each wash was followed by centrifugations at 2600 rpm for 1-2 min. For peptide purification, samples were centrifuged at 13000 rpm for 5 min and the loaded onto StageTips. Samples were centrifuged at 2600 rpm for 5 min, StageTips were washed with buffer A and centrifuged at 2 600 for 3 min. Finally, StageTips were washed twice with buffer B and each time centrifuged at 2 600 for 3 min and stored at 4°C untill submission to the CECAD proteomics facility for further processing.

Samples were analyzed by the CECAD Proteomics Facility on an Orbitrap Exploris 480 (Thermo Scientific, granted by the German Research Foundation under INST 1856/71-1 FUGG) mass spectrometer equipped with a FAIMSpro differential ion mobility device that was coupled to an UltiMate 3000 (Thermo Scientific). Samples were loaded onto a precolumn (Acclaim 5µm PepMap 300 µ Cartridge) for 2 min at 15 ul flow before being reverse flushed onto an in-house packed analytical column (30 cm length, 75 µm inner diameter, filled with 2.7 µm Poroshell EC120 C18, Agilent). Peptides were chromatographically separated at a constant flow rate of 300 nL/min and the following gradient: initial 6% B (0.1% formic acid in 80 % acetonitrile), up to 32% B in 72 min, up to 55% B within 7.0 min and up to 95% solvent B within 2.0 min, followed by column wash with 95% solvent B and re-equilibration to initial condition. The FAIMS pro was operated at -50V compensation voltage and electrode temperatures of 99.5 °C for the inner and 85 °C for the outer electrode. For the Gas-phase fractionated library, a pool generated from all samples was analyzed in six individual runs covering the range from 400 m/z to 1000 m/z in 100 m/z increments. For each run, MS1 was acquired at 60k resolution with a maximum injection time of 98 msec and an AGC target of 100%. MS2 spectra were acquired at 30k resolution with a maximum injection time of 60 msec. Spectra were acquired in staggered 4 m/z windows, resulting in nominal 2 m/z windows after deconvolution using ProteoWizard (Chambers, 2012). For the samples, MS1 scans were acquired from 399 m/z to 1001 m/z at 15k resolution. Maximum injection time was set to 22 msec and the AGC target to 100%. MS2 scans ranged from 400 m/z to 1000 m/z and were acquired at 15 k resolution with a maximum injection time of 22 ms and an AGC target of 100%. DIA scans covering the precursor range from 400 - 1000 m/z and were acquired in 60 x 10 m/z windows with an overlap of 1 m/z. All scans were stored as centroid.

The gas-phase fractionated library was built in DIA-NN 1.8.1 (Demichev 2020) using A Swissprot mouse canonical database (UP589, downloaded 04/01/22) with settings matching acquisition parameters.Samples were analyzed in DIA-NN 1.8.1 as well using the previously generated library and identical database. DIA-NN was run with the additional command line prompts “—report-lib-info” and “—relaxed-prot-inf”. Further output settings were: filtered at 0.01 FDR, N-terminal methionine excision enabled, maximum number of missed cleavages set to 1, min peptide length set to 7, max peptide length set to 30, min precursor m/z set to 400, max precursor m/z set to 1000, cysteine carbamidomethylation enabled as a fixed modification. Afterwards, DIA-NN output was further filtered on library q-value and global q-value <= 0.01 and at least two unique peptides per protein using R (4.1.3). Finally, LFQ values calculated using the DIA-NN R-package. Afterwards, analysis of results was performed in Perseus 1.6.15 (Tyanova 2016).

GO analysis of up and downregulated pathways in the cerebellum of 2 and 3 months old mice was performed using ShinyGO v0.80 (South Dakota State University; Ge, Jung and Yao, 2020). Venn diagram analysis was performed using Venny2.1 (Oliveros, J.C. (2007-2015) Venny). An interactive tool for comparing lists with Venn's diagrams. <https://bioinfogp.cnb.csic.es/tools/venny/index.html>).

**APEX proteomics**

In order to study Purkinje cell-specific proteome in the mouse cerebellum, we used an enzyme-catalyzed proximity labeling approach combined with mass spectrometry-based proteomics. For proximity labeling within genetically targeted neurons, a Cre-dependent AAV expressing the engineered ascorbate peroxidase APEX2 (ssAAV-1/2-hEF1α-DIO-dAPEX2) was intracranially injected into the cerebellum of 6 weeks old AP-2 cKO and control mice. For stereotactic surgery procedure see *Stereotactic viral injection* below. Three weeks after stereotactic surgery, injected mice were anesthetized with an intraperitoneal injection of Ketanest/Rompun and transcardially perfused with ice cold cutting solution (92 mM N-Methyl-D-glucamine, 2.5 mM KCl, 30 mM NaHCO_3_, 20 mM HEPES, 1.25 mM NaH_2_PO_4_, 2mM thiourea, 5 mM sodium ascorbate, 3 mM sodium pyruvate, 10 mM MgSO_4_, 0.5 mM CaCl_2_, 25 mM D-glucose, pH 7.4 and saturated with 95% O2/5%CO2). Afterwards, the brain was rapidly extracted, the cerebellum isolated and chopped into 300 µm pieces with a tissue chopper (Cavey Laboratory Engineering Co. LTD). Chopped cerebellar tissue was incubated in aCSF (125.2 mM NaCl, 2.5 mM KCl, 26 mM NaHCO_3_, 1.3 mM MgCl_2_ 6 H_2_O, 2.4 mM CaCl_2_, 0.3 mM NaHPO_4_, 0.3 mM KH_2_PO_4_ and 10mM D-glucose) supplemented with 0.5 mM biotinphenol (BP) at 37°C for 30 min (95%O_2_/5% CO_2_). Afterwards, APEX labeling was intiated by the addition of 1 mM H_2_O_2_ to aCSF at room temperature. After one minute, aCSF was discarded and exchanged with cold quenching buffer (aCSF supplemented with 10mM Trolox, 20 mM sodium ascorbate and 10 mM NaN_3_) on ice. The tissue was washed twice with cold quenching buffer (in-between incubation times of 2 minutes) and twice more with cold PBS. Finally, samples were resuspended in 8M urea buffer supplemented with protease inhibitor and centrifuged for 15 min at 20000g at room temperature. The supernatant was collected and transferred in fresh 1.5 ml Eppendorf tubes. The protein concentration was measured and all samples were adjusted to reach the same concentration with 8M urea. Finally, an acetone precipitation protocol was applied. Briefly, samples were mixed with 4 times the volume of cold (-20°C) acetone, vortexed and incubated at -20°C for 3 hours. Afterwards, samples were centrifuged for 10 min at 15000g at 4°C and the supernatant discarded. The protein pellet was washed twice with 80%-90% acetone (-20°C) and centrifuged for 10 min at 15000g at 4°C. The supernatant was decanted and the acetone was allowed to evaporate at room temperature for approximatively 10 min. The pellet was resuspended in 5 µL/10µg protein of 6M urea in ABC solution and afterwards sonicated. Samples were stored at -80°C until enrichment on SAV-beads and MS experiments.

Biotinylated proteins were captured using Pierce™ High Capacity Streptavidin Agarose (Thermo Scientific, #20359). Briefly, the streptavidin bead slurry was washed three times with 1x PBS. The washed beads were then incubated with the protein sample overnight at 4°C with gentle tumbling. After incubation, the beads were washed extensively with 1x PBS (five washes). Finally, the beads were resuspended in a sufficient volume of 8M Urea solution to completely cover the beads, maximizing the yield of enriched proteins. Following protein capture, samples were processed according to the established proteomics protocol provided by the CECAD proteomics facility (mentioned above).

**Adeno-associated viruses’ generation**

*DNA constructs*

pAAV/L7-6-EGFP-WPRE was generated from pAAV/L7-6-GFP-WPRE (a gift from Hirokazu Hirai, Addgene plasmid # 126462 ; http://n2t.net/addgene:126462 ; RRID:Addgene_126462) and the pAAV backbone from pAAV-hSyn-mScarlet (a gift from Karl Deisseroth, Addgene plasmid #131001; http://n2t.net/addgene:131001; RRID:Addgene_131001) using NEBuilder HiFi DNA Assembly (NEB). A P2A-Cre construct was generated from pAAV.CMV.HI.eGFP-Cre.WPRE.SV40 (a gift from James M. Wilson, Addgene plasmid # 105545 ; http://n2t.net/addgene:105545 ; RRID:Addgene_105545) by site directed mutagenesis using forward (GGCGACGTGGAGGAGAACCCCGGCCCCGCCGGCAGCATGTCCGGAGAGCAAAAGCTG) and reverse (TCTCCTCCACGTCGCCGGCCTGCTTCAGCAGGCTGAAGTTGGTGGCGCCGCTGCCCTTGTACAGCTCGTCCATGC) primers. Subsequently, pAAV/L7-6-EGFP-P2A-Cre-WPRE was generated using NEBuilder HiFi DNA Assembly. The plasmids pAdDeltaF6 (Addgene plasmid #112867; http://n2t.net/addgene:112867; RRID:Addgene_112867) and pAAV2/rh10 (Addgene plasmid # 112866 ; http://n2t.net/addgene:112866 ; RRID:Addgene_112866) were a gift from James M. Wilson. All DNA constructs were confirmed by Sanger sequencing (Eurofins).

*rAAV2/rh10 preparation*

Recombinant AAV2/rh10 particles were prepared in HEK293T cells (DSMZ no. ACC 635) by transfecting either pAAV/L7-6-EGFP-WPRE or pAAV/L7-6-EGFP-P2A-Cre-WPRE together with pAdDeltaF6 and pAAV2/rh10. Viral particles were precipitated with PEG/NaCl and cleared with chloroform extraction (Kimura et al., 2019). AAVs were purified by adapting scalable anion-exchange chromatography strategies (Dickerson, Argento, Pieracci, & Bakhshayeshi, 2021; Wang et al., 2019). Cleared AAVs were concentrated roughly 20-fold with pre-washed (PBS + 0.001% (v/v) Poloxamer 188, Sigma Aldrich) 100 kDa Amicon filters (Merck/Millipore) and diluted 10-fold in buffer A (10 mM bis-tris-propane pH 9.0, 1 mM MgCl_2_). AAVs were applied at a flow-rate of 3 mL/min to a self-packed 1 mL column (POROSTM HQ 50 µm strong anion exchange resin, Thermo Fisher Scientific), which was equilibrated in buffer A. After injection, the column was rinsed with 20 column volumes buffer A, washed with 20 column volumes 4 % buffer B (10 mM bis-tris-propane pH 9.0, 1 mM MgCl_2_, 1 M NaCl). AAVs were eluted with 35 % buffer B. Eluted fractions were concentrated and buffer exchanged to PBS + 0.001% (v/v) Poloxamer 188 using 100 kDa Amicon filters. Purity of viral preparations were assessed with SDS-PAGE/Colloidal Commassie staining and AAV titers determined using Gel green® (Biotium) (Xu, DeVries, & Zhu, 2020).

**Stereotactic viral vector injection**

Stereotactic injections of AAV1/2-Ef1α-DIO-EYFP (pAAV-Ef1a-DIO-EYFP was a gift from Karl Deisseroth, Addgene viral prep # 27056-AAV1, titer ≥ 1x10^13^ vg/mL) were performed on 4week-old AP-2 cKO mice and control littermates to label PCs with EYFP in a Cre-dependent manner. For AP-2µ acute deletion in the cerebellum, stereotactic injections of AAV2/rh10-L7-6-EFGP-WPRE control virus (titer 5.05 *10^13^ GC/mL) or AAV2/rh10-L7-6-EFGP-P2A-Cre-WPRE (titer 5.04 *10^13^ GC/mL) were performed on 6 weeks old *Ap2m1^flox/flox^* mice. Mice were weighed and anaesthetised with an intraperitoneal injection of Ketamine (100 mg/kg)/ Xylazine (20 mg/kg)/ Acepromazine (3 mg/kg) and placed into a stereotaxic apparatus (David Kopf Instruments) in absence of pedal reflexes. An eye ointment was applied on the eyes to avoid drying of the corneas and a local painkiller was subcutaneously injected before opening the skin and cleaning the skull using NaCl. The stereotactic landmark Bregma was identified with the help of a DINO Lit D-sub VGA microscope (VWR) and used to calculate the final stereotactic coordinates to target lobes IV/V of the cerebellum (AP: -5.63 mm, ML: 0 mm and DV: -1 and – 0.75 mm). A small hole was drilled using a micro drill (WPI) and 300 nl of AAV was injected using a 34 g beveled NanoFil needle (WPI), a 10 µl NanoFil syringe (WPI) and a microinjection pump (WPI) to control the injection speed (100 nl/min). After the injection at each depth, the syringe was kept in place for 3 min and then slowly retracted. Afterwards, the skull was re-hydrated with NaCl, the wound was closed, and mice were given a dose of carprofene intraperitoneally (100 µl/10g BW) to reduce postsurgical pain. A subcutaneous injection of 5% glucose solution (100 µl/10g BW) and mice were placed on a hot plate (Labotect) at 37°C was given to enhance postoperative recovery. 2 weeks (for AAV1/2-Ef1α-DIO EYFP injections) or 4 weeks (for AAV2/rh10-L7-6-EFGP-WPRE or AAV2/rh10-L7-6-EFGPP2A-Cre-WPRE injections) mice were transcardially perfused.

**Multiplex fluorescent *in situ* hybridization**

Transcardial perfusion and tissue processing was performed as described above (“Tissue processing and immunohistochemical analysis”). 40 µm sagittal cerebellar sections from 6-week-old mice previously injected with AAV1/2-Ef1α-DIO-EYFP were mounted on SuperFrost Ultra plus microscope slides (Thermo Scientific) and dried overnight at RT. Multiplex fluorescent RNA *in situ* hybridization was performed using RNAscope Fluorescent Multiplex Detection Reagents (323110, ACDBio, Newark, CA, USA) according to the instructions provided by the manufacturer for frozen tissue. The probe for Grid2ip (Mm-Grid2ip-C1) was designed by ACDBio (Cat No. 1253061-C1). Hybridized probe was detected with Opal 650 (PerkinElmer). Sections were mounted using ProLong Gold Antifade Mounting Medium with NucBlue Staining (Invitrogen), sealed with nail polish and imaged 24h later at a confocal microscope Stellaris (Leica) equipped with a 63x objective (HC PL APO 63x/1.30 GLYC CORR CS2, Stellaris).

**Behavioral tests**

*Rotarod*

2 months old AP-2 cKO mice and control littermates were acclimated to the rotarod apparatus (Ugo Basile) in a 5-minutes run on a rod rotating at a constant speed of 8 rpm for 2 days in a raw. On the third day, the mice went through 3 test runs where they had to balance on the rotating rod for 5 min while the speed increased from 4 to 40 rpm within 5 min. Between runs, a 30 min break was kept. The duration that each mouse was able to stay on the rotating rod in each run was recorded as time to fall and used as parameter for the analysis. The time to fall was then averaged across the 3 runs.

*DigiGait treadmill test*

The Digigait motorized transparent treadmill (Mouse Specifics, Inc.) was used to assess locomotor performance. Recording of animals from a ventral view was possible thanks to a high-speed video camera placed below the transparent belt of the motorized treadmill. 2 months old AP-2 cKO mice and control littermates were allowed to explore the treadmill compartment the day before the actual experiment for 1 min at the speed of 8 cm/s. The day of the experiment, the speed of the belt was increased from 8 cm/s to 14, 18, 24 and 30 cm/s for video recording depending on the locomotor capability. The percentage of mice able to perform the task at each speed for each genotype was calculated and shown in the analysis as DigiGait performance.

*SIMI Motion test*

2-month-old AP-2 cKO mice and control littermates were tested to cross beams of 1.3 meters in length and 5, 12 or 25 mm wide. Individual trials were recorded through 8 high-speed cameras (mV Blue Cougar XD; 200 frames/second) strategically positioned in a circular arrangement around the beam. The camera's positioning allowed the performance capture from 8 different angles. For detailed analysis, the camera parallel to the beam was used for the 2D reconstruction of joint kinematics. Each mouse crossed every beam for a minimum of 3 times. The number of slips per run was counted and averaged across all runs. The limb kinematics of individual step cycles were normalized to account for differences in the duration of stance and swing and averaged for each mouse and then for each genotype.

Manual annotation of the step cycle and tracking using DeepLabCut on markerless animals were utilized for kinematic analysis. The manual annotation required the frame-by-frame examination of the videos to determine the beginning of the swing phase, the end of the swing phase/start of the stance phase, and the end of the stance phase. Steps resulting in footslips were excluded from the kinematic analysis, and individual steps were demarked from swing (forward propulsion of the limb) to stance (support phase) phase. Only 2 AP-2 cKO mice were analysed for the 5 mm beam since the remaining animals were not able to cross the beam. We did not exclude any of the control mice for the narrower beam. DeepLabCut was used to track the 2D coordinates of the mice and to track the position of the beam in the video and establish the baseline for the vertical axis. The coordinates extracted by DeepLabCut were uploaded to AutoGaitA (Hosseini et al., bioRxiv 2024) to calculate angles and velocities, to normalize the coordinate across step cycles, to average trials per mouse and to average groups per genotype. The grouped values were analyzed using GraphPad Prism version 9.5.1 (GraphPad Software, Inc., USA).

**Echo-MRI body composition analysis**

The body composition of 2 months old AP-2 cKO male mice and control littermates was measured using an EchoMRI-100H Body Composition Analyzer (EchoMRI ®). The whole-body masses of fat, lean, free water and total water were measured. Mice were placed in an animal holder and measures were taken 3 times per mouse, for a duration of 0.5-3.2 min each. The measurements were then averaged between the 3 runs and shown as percentage of body weight.

**In vitro whole cell recordings**

1-month-old AP-2 cKO mice and control littermates were used for recordings in acute brain slices. Animals were decapitated after being deeply anesthetized by isoflurane inhalation overdose. The brain was rapidly placed in ice-cold artificial cerebrospinal fluid (ACSF, in mM: 124 NaCl, 2.5 KCl, 1.25 NaH_2_PO_4_, 26 NaHCO_3_, 1 MgSO_4_, 2 CaCl_2_, 25 D-Glucose and 1mM kynurenic acid), bubbled with 95%/5% O2/CO2 (pH 7.4). Acute coronal 250 µm slices containing the cerebellum were prepared with a vibratome (Campden Instruments 7000smz-2), placed in warm ACSF without kynurenic acid bubbled with 95%/5% O2/CO2 (pH 7.4), and maintained at 32-34°C for ~30 min, then cooled to room temperature (22-24°C) for at least one hour before use. For experiments, slices were transferred to the recording chamber and superfused (2 ml.min-1) with ACSF without kynurenic acid at 32°C bubbled with 95%/5% O2/CO2 (pH 7.4). Purkinje neurons were identified with an Olympus 40 x water-immersion objective. Pipettes with resistance 5-6 MΩ made of borosilicate glass capillaries with O.D. 1.5 mm, I.D. 0.86 mm (Sutter, Item# BF-150-86-10) were prepared using P-97 micropipette puller (Sutter Instruments, Item# P-97) and contained (in mM): 127 K-gluconate, 8 KCl, 10 phosphocreatine, 10 HEPES, 4 Mg-ATP, 0.3 Na-GTP (osmolality, 285 mOsm; pH 7.2 adjusted with KOH). Somatic whole-cell current-clamp recordings were made from cerebellar Purkinje neurons using Multiclamp 700B amplifier and Digidata 1550B digitizer (Molecular Devices). Data were acquired with Clampex 11.2 (Molecular Devices), digitized at 10 kHz. Electrophysiology data analysis was performed using Clampfit 11.3 (Molecular Devices).

**Cerebellar acute slices**

1 month old AP-2 cKO mice and control littermates were sacrificed via cervical dislocation. Brains were isolated and cerebellum dissected. 200 µm horizontal acute slices were obtained from the cerebellum with a VT1200 Vibratome (Leica). While cutting, the cerebellum was submerged in ice-cold, carbogen saturated (95% O_2_ and 5% CO_2_) low-Ca^2+^ artificial cerebrospinal fluid (ACSF: 125 mM NaCl, 2.5 mM KCl, 1.25 mM sodium phosphate buffer 0.4 M, 25 mM NaHCO_3,_ 25 mM glucose, 0.5 mM CaCl_2_ and 3.5 mM MgCl_2,_ osmolarity adjusted between 310 and 330 milliosmoles, pH = 7.4).

**MG132 experiments**

To test whether GRID2IP levels were changed after proteasome inhibition, cerebellar acute slices from 1-month-old AP-2 cKO and control littermates were treated with the proteasome inhibitor MG132. Slices were incubated for 6.5h at 37°C/5% CO_2_ with 100 µM MG132 or 100 µM DMSO (control) diluted in culturing medium (MEM, 0.00125% ascorbic acid, 10 mM D-glucose, 1 mM GlutaMAX^TM^; 20% (v/v) horse serum, 0.01 mg/ml insulin, 14.4 mM NaCl; 1% P/S). Afterwards, slices were fixed for 1h at RT with 4% PFA before immunohistochemistry was performed.

**Transferrin uptake assay in cerebellar acute slices**

For transferrin (Tfn) uptake, 1 month old AP-2 cKO mice and control littermates were sacrificed via cervical dislocation. Brains were isolated and cerebellum dissected. 100 µm horizontal acute slices were obtained from the cerebellum with a VT1200 Vibratome (Leica). While cutting, the cerebellum was submerged in ice-cold, carbogen saturated (95% O_2_ and 5% CO_2_) low-Ca^2+^ artificial cerebrospinal fluid (ACSF: 125 mM NaCl, 2.5 mM KCl, 1.25 mM sodium phosphate buffer 0.4 M, 25 mM NaHCO_3,_ 25 mM glucose, 0.5 mM CaCl_2_ and 3.5 mM MgCl_2,_ osmolarity adjusted between 310 and 330 milliosmoles, pH = 7.4). Slices were incubated in ACSF for 2 h at 37°C/5% CO_2_ before incubation for 1 h with 25 µg/µl human Tfn conjugated to Alexa Fluor 488 (Invitrogen) in ACSF. Cell surface-bound Tfn was removed by an ice-cold acid wash (0.2 M acetic acid + 0.5 M NaCl, pH 2.8) for 5 min then rinsed with ice-cold PBS 3 times. Afterwards, slices were fixed for 1h at RT with 4% PFA before immunohistochemistry was performed.

**Primary cerebellar culture**

*Culture preparation*

Postnatal day 8 (P8) AP-2 cKO pups and control littermates were decapitated and brains were collected in ice cold solution B (300 mg BSA (fatty acid free), 1.5mM MgSO4 and 13 mM glucose in 100 ml dPBS). The cerebellum was isolated and chopped into 700 µm thick pieces with a tissue chopper (Cavey Laboratory Engineering Co. LTD). The chopped tissue was subsequently incubated in solution T (solution B plus 1mg/4mL trypsin) at 37°C/5% CO_2_ for 15 min. In order to stop trypsinization, solution C (600U DNase; 0.5 mg Soybean Trypsin Inhibitor (SBTI); 100 μl MgSO_4_ in 10 ml solution B) was added to the samples (twice the volume as solution T), which were then centrifuged at 1000g for 1 min at 4°C. The supernatant was removed and cell suspension was dissolved in 1 mL growth medium (5 mL FBS in 50 ml stock medium: 500 ml DMEM; 2 ml B-27, 0.74 g KCl, 10 mM glucose, 1 mM sodium pyruvate, 5 ml P/S). Afterwards, the cell suspension was mechanically dissociated with a fire-polished glass pipette. Dissociated cells were layered on top of 5 mL EBSS (EBSS plus % (w/v) bovine serum albumin (BSA); 3 mM 3.82% MgSO4) and centrifuged at 1500 g for 5 min at 4°C. Afterwards, the supernatant was discarded and the pellet dissolved in growth medium and cell density determined using a Neubauer counting chamber. Finally, cells were plated at a density of 750000 cells per coverslip and fed after one hour with growth medium. After 24h, half of the medium was replaced with fresh growth medium plus 4 µM AraC. Cerebellar neurons were cultured for 19 days (DIV 19) under constant conditions at 37°C/5% CO_2_.

*Primary cerebellar culture viral transduction*

When necessary, in order to identify PCs in primary cerebellar cultures, neurons were transduced on DIV3 with AAV2/rh10-L7-6-EFGP-WPRE (0.3 µl/1mL of medium; titer 5.05 *10^13^ GC/mL).

PitStop2-mediated CME inhibition

On DIV18, primary cerebellar cultures from WT mice were incubated with 30µM PitStop2 (Abcam) or DMSO (control) diluted into cerebellum growth medium at 37°C/5% CO_2_ for 24h. Afterwards, neurons were fixed at DIV19 with 4% PFA for 10 min at RT and used for experiments.

Transferrin uptake assay

For transferrin uptake, cerebellar growth medium was removed from DIV19 primary cerebellar cultures from AP-2 cKO mice and control littermates (previously transduced with AAV2/rh10-L7-6-EFGP-WPRE) and replaced with pre-warmed Neurobasal-A Medium (Thermo Fisher Scientific). Neurons were starved in this medium for 2h at 37°C/5% and then incubated for 30 min at 37°C/5% with the same medium containing 15 μg/ml human Transferrin conjugated to Alexa Fluor 568 (BioTrend). Cell-surface bound Transferrin was removed by 3 PBS washes and immediately fixed with 4% PFA for 10 min at RT.

Transferrin uptake was also used to prove CME inhibition by Pitstop2 in WT primary cerebellar culture. For this purpose, DIV18 neurons were incubated with 30µM Pitstop2 (Abcam) or DMSO (control) diluted into cerebellum growth medium at 37°C/5% CO_2_ for 24h. On DIV19, neurons were starved for 2h at 37°C/5% in a pre-warmed Neurobasal-A medium containing 30µM Pitstop2. Afterwards, 15 μg/ml human Transferrin conjugated to Alexa Fluor 488 were added to the medium for 30 min at 37°C/5%. Cell-surface bound Transferrin was removed by 3 PBS washes and immediately fixed with 4% PFA for 10 min at RT.

Immunocytochemistry

Fixed primary cerebellar cultures were blocked for 1h at RT in 0.3% saponin (Sigma) in PBS plus 5% NGS. Afterwards, cells were incubated for 1h at RT with primary antibodies (see Key Resource Table) diluted in 0.3% saponin/PBS plus 5% NGS. Cells were washed 3 times with PBS and then incubated for 30 min at RT with secondary antibodies (see Key Resource Table) diluted in 0.3% saponin/PBS plus 5% NGS. Finally, cells were washed 3 times with PBS and coverslips were mounted on glass slides with Immu-Mount (Epredia).

**Cerebellar organotypic cultures (OTCs)**

OTCs preparation

P8 AP-2 cKO pups and control littermates were decapitated and brains were collected in ice-cold HBSS (Gibco). Cerebellum was isolated and 300 µm sagittal slices were obtained with a tissue chopper (Cavey Laboratory Engineering Co. LTD). Sections were collected and washed 3 times by being carefully transferred with a glass pipette in 3 separate dishes containing pre-warmed HBSS. Afterwards, slices were transferred onto Millicell Standig Cell Culture Inserts (Merk Millipore) and fed with culturing medium (MEM, 0.00125% ascorbic acid, 10 mM D-glucose, 1 mM GlutaMAX^TM^, 20% (v/v) horse serum, 0.01 mg/ml insulin, 14.4 mM NaCl, 1% P/S) from below the cell culture insert membrane. Every 2 days, the medium was replaced and slices were cultured for 21 days under constant conditions at 37°C/5% CO_2_.

*OTCs viral transduction*

OTCs were used for calcium imaging. Therefore, slices were transduced at DIV1 by applying 1 µl of ssAAV-9/2-mCaMKIIα-jGCaMP7f-WPRE-bGHp(A) (titer 1.1x10^13^, Viral vector facility Zurich) on top of each slice. Slices were cultured until DIV21 at 37°C/5% CO_2_ until they were used for experiments.

Ceftriaxone treatment

Ceftriaxone enhances the expression of excitatory amino acid transporter 2 (EAAT2/ GLT-1), one of the major glutamate transporters primarily expressed in astrocytes. Ceftriaxone has therefore the potential to manipulate glutamate transmission and ameliorate neurotoxicity. OTCs from AP-2 cKO mice and control littermates previously transduced with ssAAV-9/2-mCaMKIIα-jGCaMP7f-WPRE-bGHp(A) were cultured for 15 days in OTC culturing medium at 37°C/5% CO_2._ On DIV 15, slices were treated with 100 µM Ceftriaxone for 7 consecutive days until DIV21. On DIV21, OTCs were either used for calcium imaging experiments or shock-frozen in liquid nitrogen and stored at -80°C until tissue lysis for immunoblotting analysis. Untreated OTCs were used as control.

**Calcium imaging experiments**

DIV 21 OTCs from AP-2 cKO mice and control littermates were used for GCaMP7f-based calcium imaging. Slices were cut out from the cell culture inserts and placed into a RC-47FSLP stimulation chamber (Warner Instruments) filled with OTC imaging medium (2 mM CaCl_2_, 10 mM D-glucose, 3 mM KCl, 1 mM MgCl_2_, 136 mM NaCl, 24 mM NaHCO_3_, 1.25 mM NaH_2_HPO_4_ in dH_2_O) bubbled with 95%/5% O_2_/CO_2_ and constantly perfused. A Multiphoton microscope SP8 (Leica) equipped with a 20x/0.75 multi-immersion objective was used for confocal live imaging. 60s recordings (1 to 3 recording per slice) were acquired at 512x512 pixel resolution, bidirectional at 1 frame per second. After 20s of baseline recording, slices were stimulated once with a 100 Hz pulse and recorded for 40 more seconds, for a total of 60s for each recording.

**Calcium imaging analysis**

*Cell/dendritic patches detection and signal extraction*

Signal extraction from calcium imaging recordings was done using ImageJ (Fiji). Region of interests (ROIs) were drawn around PCs cell bodies or dendritic patches identified thanks to GFP expression driven by the ssAAV-9/2-mCaMKIIα-jGCaMP7f-WPRE-bGHp(A) virus. The mean grey value from each ROI was extracted for every single frame (60 in total) and used for the analysis.

*Automatic cell curation and filtering*

Ca^2+^ signals were sampled at a rate of 1 Hz. Individual Ca^2+^transients were normalised to the 5 s-period preceding electrical stimulation. To identify low quality components, we performed k-means clustering (k=5, nsets=20, kmeans function, stats package, R). The number of clusters was optimised according to the total within-cluster sum of squared distances. We identified a distinct cluster of low-quality components with continuous baseline shifts, indicative of motion artefacts or bleaching. Components in this cluster were discarded from further analysis (11-19 % of the total). All other components were used for further analysis.

*Characterization of stimulus responses*

*Heatmaps.* For visual inspection of Ca^2+^ transients in the form of activity maps, individual transients were smoothed (Tukey’s smoothing with running median of 3, smooth function, stats package, R), sorted according to the median signal amplitude of the 10 s-period following stimulation.

*Individual response parameters.* The following parameters were used to assess stimulus responses of individual components:

| **Parameter** | **Description** |
| --- | --- |
| Average Ca^2+^ response | Arithmetic mean of the Ca^2+^ signal within the 10 s-period following electrical stimulation |
| Peak Ca^2+^ response | Highest Ca^2+^ signal within the 40 s-period following electrical stimulation |
| Time to peak | Time point of the maximum Ca^2+^ response within the 40 s-period following electrical stimulation |

*Synchronicity*. To assess the degree of similarity in firing patterns of individual Ca^2+^ transients, we calculated the pair-wise Pearson correlation coefficient for each component with all other components within the 40 s-period following electrical stimulation per genotype and condition. For visual inspection, the correlation matrix per genotype and condition was sorted according to hierarchical cluster analysis (cor_sort function, correlation package, and hclust function, stats package, R).

**Image acquisition**

Images of fluorescently stained brain slices and primary cerebellar cultures were acquired at a confocal microscope Stellaris (Leica) or at a confocal microscope SP8 (Leica). At 20x (HC PL APO 20x/0.75 CS2, Stellaris) magnification, single plane tile images were acquired for stained brain slices. At 40x (HC PL APO 40x/0.95 CORR, Stellaris; PL Apo 40x/0.85 CORR CS, SP8) and 63x (HC PL APO 63x/1.30 GLYC CORR CS2, Stellaris; PL Apo 63x/1.40 Oil CS2, SP8) either single plane images or z-stacks were acquired for both stained brain slices and primary neurons. Z-stacks are shown as maximum intensity projection images. Tile images were merged with LAS X software (Leica).

*In situ* hybridized sections were imaged at a confocal microscope Stellaris (Leica) equipped with a 63x objective (HC PL APO 63x/1.30 GLYC CORR CS2, Stellaris). Z-stacks were acquired and images are shown as maximum intensity projection of z-stack.

Brightfield images were acquired with a slide scanner microscope (S360 Hamamatsu) equipped with a 40x objective.

**Analysis of fluorescently stained brain slices**

The following analysis was used to quantify GRID2IP levels in 200 µm thick cerebellar sagittal acute slices treated with MG132 or DMSO,GLURδ2 levels in distal dendrites and and AP-2α in PC somata in 40 µm thick sagittal cerebellar slices as well as transferrin-uptake in 100 µm thick cerebellar sagittal acute slices. The mean grey value of the GRID2IP, AP-2α staining or transferrin signal in PCs somata or GLURδ2 in distal dendrites was extracted with ImageJ (Fiji) and normalized for background fluorescence (area devoid of GRID2IP,GLURδ2, AP-2α or transferrin-488 signal).

**Analysis of in situ hybridized sections**

For analysis of *Grid2ip* mRNA puncta, maximum intensity projections of acquired z stack were generated for both RNAscope probe and GFP (AAV1/2-Ef1α-DIO EYFP) channels using ImageJ (Fiji). The GFP fluorescence mediated by the AAV transduction of PC was used to draw ROIs including both PCs somata and dendrites. These same ROIs were copied onto the *Grid2ip* channel and allowed to mark the region where RNA puncta were quantified. For this purpose, a threshold was set on the channel showing *Grid2ip* mRNA expression, allowing to label exclusively the signal coming from mRNA puncta. Puncta where finally quantified using the particle analysis tool in Fiji and the total number of puncta was normalized by the ROI area for each single image.

**Cell number quantification**

Cresyl-violet stained sagittal cerebellar sections from 1-, 2- and 3-month-old mice were used to quantify the number of PCs in AP-2 cKO mice and control littermates. Cerebellar lobes from lobe I to lobe X were identified in 3 to 5 sections from each mouse, ranging from Bregma 0.48 mm to Bregma 1.2 mm according to the Allen Brain Atlas. PCs number was counted in each lobe for each Bregma coordinate and then averaged, in order to obtain the total number of neurons per lobe. Aperio Image Scope image viewing software (Leica) was used to analyze acquired images.

**Analysis of perisomatic puncta**

Perisomatic quantification of vGLUT2 and vGLUT1 puncta was performed on fluorescently immunostained sagittal cerebellar sections from WT and AP-2 cKO 1- and 3-month-old animals. Z-stack confocal images were acquired at a 63x magnification and maximum intensity projections extracted from these stacks were used for this analysis. Using ImageJ, the GFP fluorescence mediated by the AAV transduction or the tdTomato fluorescence of PCs was used to draw ROIs around somata. The number of puncta per cell was quantified and normalized to the area of the soma and used for further statistical analysis.

**Colocalization analysis**

Colocalization analysis between AP-2α and GRID2IP was performed on fluorescently immunostained sagittal cerebellar sections from 2-month-old WT mice. Non-processed raw dual channel images were used for the analysis. ROIs were manually drawn in order to outline somata, primary or secondary dendrites. Pearson´s correlation coefficient (Rp) was determined using the JACoP plug-in in ImageJ. The same analysis was performed to investigate differences in colocalization between GLURδ2 and VGLUT1 in distal PCs dendrites of 1 month old AP-2 cko mice and control littermates.

**Analysis of number of spines**

The number of spines in proximal and distal dendrites was quantified in sagittal cerebellar sections from 6 weeks old AP-2 cKO mice and control littermates previously injected with AAV1/2-Ef1α-DIO EYFP for single PCs visualization and counterstained with GFP. Z-stack confocal images were acquired at a 63x magnification and single planes extracted from these stacks were used for this analysis. Using ImageJ, a 10 µm line was drawn and the number of spines was counted within this length. 4 to 7 PCs per mouse were analyzed. For each cell, 3 different measurements were obtained and then averaged for final analysis. The same quantification was performed to determine differences in number of spines in PCs from 10 weeks old AP-2m1^flox/flox^ mice previously injected with AAV2/rh10-L7-6-EFGP-WPRE control virus or AAV2/rh10-L7-6-EFGP-P2A-Cre-WPRE virus to obtain AP-2µ acute deletion in the adult cerebellum.

**AMIRA-based 3D reconstruction and analysis**

40 µm sagittal cerebellar sections from 6 weeks old AP-2 cKO mice and littermates previously injected with AAV1/2-Ef1α-DIO EYFP for single PCs visualization and stained for GFP and GRID2IP/VGLUT1/VGLUT2 were used for the analysis. Images were acquired using HC PL APO 63x/1.30 GLYC CORR CS2 objective at a resolution of 1024 x 1024 pixels in sequential scanning frame‐by‐frame mode. Stacks of 50-80 optical sections were acquired, with a fixed stack size of 0.33 μm. 3D reconstructions were generated with Amira Software 2020.2 (Thermo Fisher Scientific). First, the surface area of single GFP-positive PCs (soma and dendrites) was reconstructed using the Amira segmentation editor. GRID2IP/VGLUT1/VGLUT2 signal was defined by generating the isosurface. Afterwards, the surface of ‘300‐nm‐distant’ GRID2IP/VGLUT1/VGLUT2‐positive voxels was mapped onto GFP‐positive reconstructed cell using the “surface distance” tool and extracted as a histogram. Values extracted from the histogram (number of voxels/μm^2^) were plotted and used for colocalization analysis.

**Statistical analyses**

Statistical analyses were conducted on mice values or cell values (indicated by data points) from at least a group of 3 mice per genotype (indicated by “N”, biological replicates) or at least 3 independent experiments (indicated by “n”, biological replicates), if not stated otherwise in the figure legend. MS Excel (Microsoft, USA) and GraphPad Prism version 9.5.1 (GraphPad Software, Inc., USA) were used for statistical analysis and result illustration (unless otherwise stated). Unpaired t-test or Welch´s unpaired t-test were used to compare the means of two groups. Statistical analysis of normalized data between the two groups was performed using a one-tailed unpaired Student´s t-test. Two‐tailed Mann‐Whitney test was used for analysis between two groups for non‐normally distributed non‐normalized data (i.e. number of spikes evoked by intracellular current injection). Statistical difference between more than two groups were compared with one-way ANOVA (Tukey’s posthoc test for multiple comparison was used to determine the statistical significance between the groups). Statistical difference between more than two groups and two conditions was evaluated using two-way ANOVA with Šidák post-hoc test. Significant differences were accepted at *P* ≤ 0.05 indicated by asterisks: **P* ≤0.05; ***P* ≤0.01; ****P* ≤ 0.001 and *****P*≤0.0001.

**Supplemental videos.**

**Videos S1,2.**

Video recordings of 2-month-old WT (Video S1) and AP-2 cKO (Video S2) mice in the DigiGait running chamber set at 24cm/sec speed, taken by a high-speed video camera mounted below a transparent treadmill belt.

**Videos S3,4.**

Kinematic analysis of 2-month-old WT (Video S3) and AP-2 KO (Video S4) mice while crossing a 25 mm beam.

**Videos S5,6.**

Kinematic analysis of 2-month-old WT (Video S5) and AP-2 KO (Video S6) mice while crossing a 12 mm beam.

**Videos S7,8.**

Kinematic analysis of 2-month-old WT (Video S7) and AP-2 KO (Video S8) mice while crossing a 5 mm beam.

**Supplemental tables.**

**Table S1**: Proteome data set from 1-month and 2-month old WT and AP-2 cKO cerebellum.

**Table S2:** Proteome data set from 6-week-old ssAAV-1/2-hEF1α-DIO-dAPEX2- injected *Ap2m1*^wt/wt^;L7*^Cre^* (WT) and *Ap2m1^fl/fl^*; L7*^Cre^* (AP-2 cKO) mice.

**Table S3**: Proteome date set of AP-2α pull-down analysis from 8-week-old WT cerebellum.

**Table S4**: Details of two-way ANOVA multiple comparison analysis of kinematic data shown in Fig. 1K-R and Fig. S2B-S.

**Table S4**: Data of patch-clamp recordings shown in Fig. 7A-C and Fig. S6A-C.
